# Oestrogen promotes innate immune evasion of *Candida albicans* through inactivation of the alternative complement system

**DOI:** 10.1101/2020.07.22.207191

**Authors:** Pizga Kumwenda, Fabien Cottier, Ben Keevan, Hannah Gallagher, Hung-Ji Tsai, Rebecca A. Hall

## Abstract

Gender is a risk factor for several infections that, for many pathogens, has been linked to sex hormones impacting host immunity and directly affecting microbial virulence. *Candida albicans* is a commensal of the urogenital tract and the predominant cause of vulvovaginal candidiasis (VVC). Factors that increase circulatory oestrogen levels like pregnancy, the use of oral contraceptives, and hormone replacement therapy predispose women to VVC, but the reasons for this are largely unknown. Here, we investigate how adaptation of *C. albicans* to oestrogen impacts the fungal host-pathogen interaction. Physiologically relevant concentrations of oestrogen promoted fungal virulence by enabling *C. albicans* to avoid the actions of the innate immune system. Oestrogen-induced innate immune evasion was mediated via inhibition of opsonophagocytosis through enhanced acquisition of the human complement regulatory protein, Factor H, on the fungal cell surface. Oestrogen induced accumulation of Factor H was dependent on the fungal cell surface protein Gpd2, with oestrogen dependent derepression of *GPD2* being mediated via a non-canonical signalling pathway involving Ebp1 and Bcr1. Therefore, we propose that, in addition to affecting the antifungal potential of vaginal epithelial cells, elevated oestrogen levels predispose women to VVC by directly enhancing fungal pathogenicity through the inactivation of complement. The discovery of this new hormone sensing pathway might pave the way in explaining gender biases associated with fungal infections and may provide an alternative approach to improving women’s health.

## Introduction

Microbial infections exhibiting gender bias are common. This bias may predispose one sex to infection over the other, or result in one sex exhibiting severer infection outcomes (Ghazeeri *et al*. 2011, García-Gómez *et al*. 2013, van Lunzen and Altfeld 2014). Sex hormones like oestrogen, testosterone and progesterone regulate many functions of the immune system, and generally males are more prone to infection than females, as overall immune responses are lower in the male population (Klein 2000, McClelland and Smith 2011, Taneja 2018). Besides the impact of sex hormones on immune cell function, sex hormones also have a direct effect on microbial pathogenicity increasing microbial persistence, metabolism and virulence gene expression (Amirshahi *et al*. 2011, Chotirmall *et al*. 2012).

The opportunistic fungal pathogen *Candida albicans* is the predominant fungal coloniser of the female reproductive tract and the major cause of genital thrush (vulvovaginal candidiasis, VVC). Approximately 75% of the female population will encounter at least one episode of VVC in their life-time, while up to 15% experience recurrent infection (RVVC) defined as four or more episodes in a twelve-month period (Sobel 1997). Although not life threatening, mucosal infections are expensive to treat, impact quality of life, and increase population morbidity.

One of the major risk factors associated with the development of VVC is elevated levels of oestrogen which occur as a result of pregnancy, the use of high oestrogen-containing oral contraceptives and hormone replacement therapy (SOBEL 1988, Dennerstein and Ellis 2001). Therefore, oestrogen plays a key role in predisposing women to VVC, but the precise mechanism(s) for this are unknown. Oestrogen promotes glycogen production at the vaginal mucosa providing a nutrient rich environment for the expansion of *C. albicans* (Dennerstein and Ellis 2001). In mouse models of VVC, were pseudoestus is induced to maintain fungal colonisation, vaginal epithelial cells have a diminished ability to control the growth of *C. albicans* (Fidel *et al*. 2000). In addition, oestrogen decreases the infiltration of phagocytes and suppresses cell mediated immunity (Salinas-Muñoz *et al*. 2018). However, in rats, preincubation of *C. albicans* with oestrogen prior to vaginal infection enhances fungal survival (Kinsman and Collard 1986), suggesting that in addition to affecting host immunity, oestrogen may directly affect the virulence of *C. albicans*. In agreement with this, oestrogen has been shown to promote hyphal morphogenesis of *C. albicans* (White and Larsen 1997), a key virulence factor of the fungus. Furthermore, an oestrogen binding protein (Ebp1) has been identified in *C. albicans* (Madani *et al*. 1994), although the importance of this protein in VVC is not known. Therefore, how *C. albicans* adapts to oestrogen is still unclear.

The *C. albicans* cell wall is a multi-layered structure consisting of an inner layer of chitin and beta-glucan, and an outer layer of heavily glycosylated proteins (Netea *et al*. 2008). The fungal cell wall is a highly dynamic structure providing rigidity, strength and protection from the environment. In addition, many components of the cell wall act as pathogen associated molecular patterns (PAMPs) and are recognised by the innate immune system (Netea *et al*. 2008, Hall and Gow 2013). Recently, remodelling of the cell wall in response to adaptation to host environments has been shown to regulate the host-pathogen interaction (Wheeler and Fink 2006, Wheeler *et al*. 2008, Hall 2015, Ballou *et al*. 2016, Hopke *et al*. 2016, Sherrington *et al*. 2017, Lopes *et al*. 2018, Pericolini *et al*. 2018, Pradhan *et al*. 2018, Pradhan *et al*. 2019, Tripathi *et al*. 2020, Williams and Lorenz 2020). In addition to direct recognition of cell wall PAMPs mediating phagocytosis, *C. albicans* also activates the alternative complement system, resulting in the deposition of complement proteins (i.e. C3) on the fungal cell surface, resulting in opsonophagocytosis (Kozel *et al*. 1996). However, *C. albicans* can evade opsonophagocytosis through the binding of complement regulatory proteins to its cell surface that inactivate the complement cascade (Meri *et al*. 2002). Here we show that *C. albicans* does adapt to oestrogen, and that this adaptation perturbs the host-pathogen interaction, inhibiting phagocytosis of the fungal pathogen. Avoidance of the innate immune system was mediated via the fungal cell surface protein, Gpd2, recruiting the human complement regulatory protein, Factor H, to the fungal cell surface inactivating the alternative complement system.

## Results

### Adaptation of *C. albicans* to oestrogen promotes innate immune evasion

Pregnant women and women taking high oestrogen-containing oral contraceptives are more prone to vulvovaginal candidiasis (VVC) (Gonçalves *et al*. 2016), while VVC occurs less frequently in postmenopausal women, indicating that oestrogen may play a role in promoting the virulence of *C. albicans*. There are four main forms of oestrogen, estrone (E1), 17β-estradiol (E2), estriol (E3) and 17α-ethynylestradiol (EE2). Estrone is the weakest form of oestrogen produced by the ovaries and adipose tissue, and is only found in menopausal women, while 17β-estradiol is the strongest form of oestrogen produced by the ovaries and has been associated with many gynaecological disorders. Estriol is a by-product from the metabolism of estradiol, and as such is found in high concentrations during pregnancy. Finally, 17α-ethynylestradiol is a synthetic oestrogen used in oral contraceptive pills. To ascertain whether adaptation of *C. albicans* to oestrogen affects the host-pathogen interaction, *C. albicans* cells were grown in the presence of physiological (0.0001 μM) and super-physiological (0.1 μM, 10 μM) concentrations of 17β-estradiol, estriol, or 17α-ethynylestradiol, and phagocytosis rates quantified. Physiological and super-physiological concentrations of all three forms of oestrogen significantly inhibited macrophage and neutrophil phagocytosis of *C. albicans*, resulting in a 50% drop in phagocytosis compared to the ethanol vehicle control (Fig 1A-C, Fig S1A, B). The reduced rates of *C. albicans* phagocytosis were independent of any impact of oestrogen on fungal growth (Fig S1C) or morphology (Fig S1D). Given that all tested forms of oestrogen elicited similar results, all subsequent experiments were performed only with 17β-estradiol.

**Figure 1.**
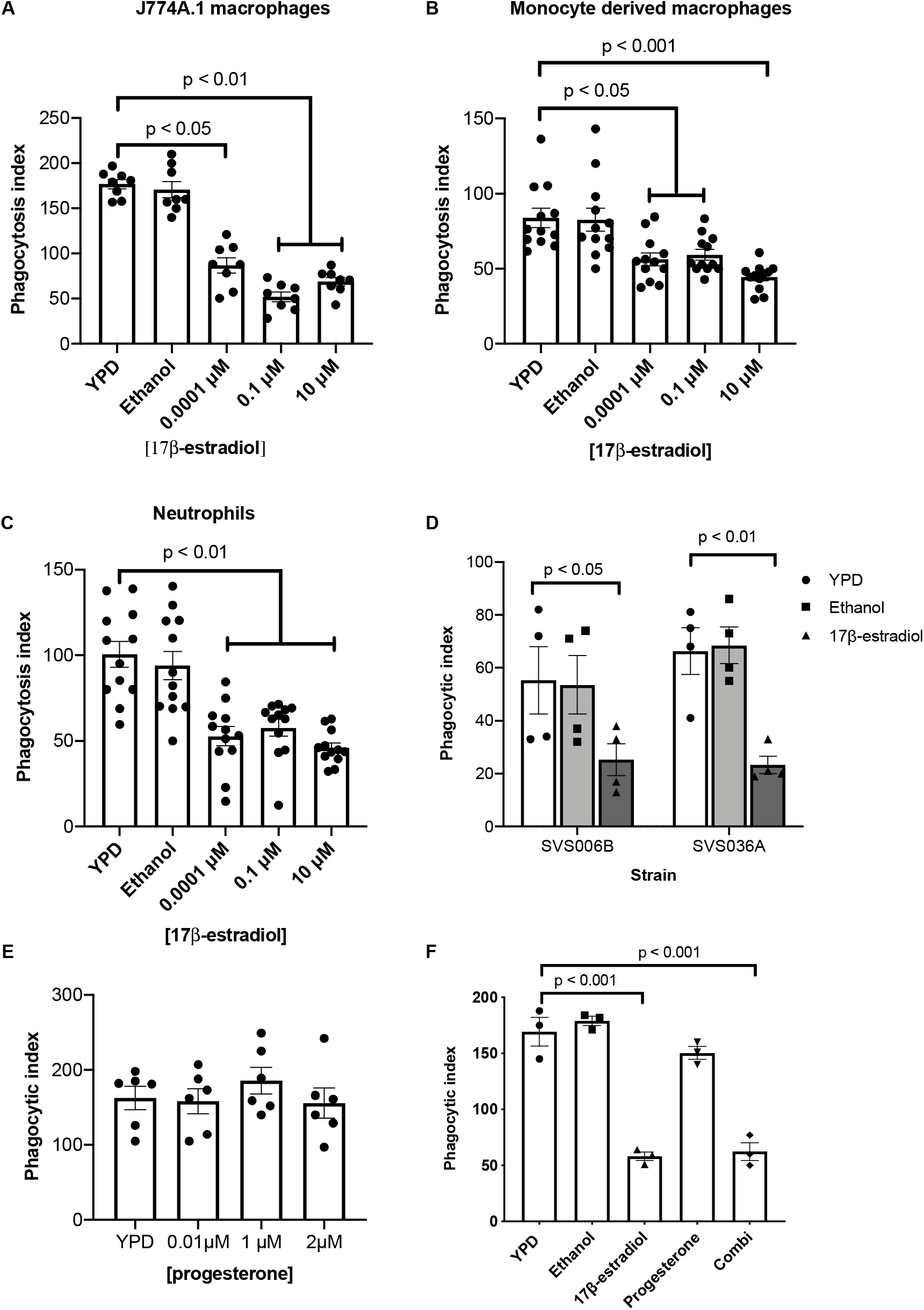
Oestrogen promotes innate immune evasion of *C. albicans*. C. albicans cells were grown in YPD with or without oestrogen supplementation. Cells were harvested, washed in PBS and co-incubated with (**A**) J774A.1 cells (**B**) human macrophages (**C**), human neutrophils. (**D**) J774A.1 phagocytosis rates of *C. albicans* clinical isolates after exposure to 10 μM 17β-estradiol. **(E)** J774A.1 phagocytosis rates of *C. albicans* after exposure to progesterone **(F)** J774A.1 phagocytosis rates of *C. albicans* after exposure to 1 μM progesterone and 10 μM 17β-estradiol, either individually or in combination (combi). Cells were fixed with 4% PFA, imaged by microscopy, scored using ImageJ software and phagocytic index determined. All data represent the mean ± SEM from at least three independent biological experiments (also see Figure S1).

To determine whether the observed inhibition of phagocytosis is truly mediated by adaptation of *C. albicans* to oestrogen, and not a result of residual oestrogen coating the surface of *C. albicans* inhibiting phagocyte function, macrophages were pre-treated with oestrogen prior to the addition of *C. albicans* or latex beads. Macrophages pre-incubated with oestrogen phagocytosed *C. albicans* at rates comparable to non-treated cells (Fig S1E), suggesting that under the tested conditions oestrogen does not directly impact the ability of macrophages to phagocytose target particles. Pre-incubating inert particles like latex beads with oestrogen had no effect on the ability of macrophages to phagocytose the particles (Fig S1F), suggesting that any residual oestrogen on the surface of particles does not interfere with the ability of macrophages to phagocytose them. In agreement with this, incubation of *C. albicans* with oestrogen on ice, which inhibits fungal growth and metabolism, did not affect phagocytosis rates (Fig S1G). Furthermore, re-inoculation of oestrogen-adapted *C. albicans* cells into fresh YPD media restored *C. albicans* phagocytosis rates (Fig S1H). Therefore, in response to oestrogen, *C. albicans* undergoes some form of adaptation that alters the host-pathogen interaction.

Finally, we tested several vaginal *C. albicans* isolates and observed that all isolates displayed reduced phagocytosis after adaptation to oestrogen, confirming that oestrogen-induced immune evasion is a general trait of clinically relevant *C. albicans* (Fig 1D). Taking this data together, we conclude that the reduction in *C. albicans* phagocytosis is due to the fungus reversibly adapting to the oestrogen, and that this adaptation interferes with the host mechanism of fungal clearance.

### Hormone induced innate immune evasion is specific to oestrogen

Progesterone been shown to affect phagocytosis rates (György 2017), and is structurally quite similar to oestrogen. Therefore, we hypothesised that *C. albicans* may adapt to progesterone in a similar way to oestrogen to evade the innate immune system. *C. albicans* cells pre-exposed to physiological and super-physiological concentrations of progesterone exhibited comparable phagocytosis rates as *C. albicans* cells grown in YPD (Fig 1E), while *C. albicans* treated with both oestrogen and progesterone still exhibited innate immune evasion (Fig 1F). Therefore, the promotion of innate immune evasion appears to be a specific and dominant attribute of oestrogen.

### Oestrogen-induced innate immune evasion is mediated via inhibition of opsonophagocytosis

The interaction between *C. albicans* and innate immune cells can be mediated via direct detection of fungal cell wall carbohydrates and opsin mediated phagocytosis (opsonophagocytosis). Adaptation of *C. albicans* to a variety of environmental conditions influences the *Candida* host-pathogen interaction, through altered Dectin-1 dependent recognition of β-glucan, which correlates with altered pro-inflammatory cytokine secretion (Wheeler and Fink 2006, Wheeler *et al*. 2008, Hall 2015, Ballou *et al*. 2016, Hopke *et al*. 2016, Sherrington *et al*. 2017, Lopes *et al*. 2018, Pericolini *et al*. 2018, Pradhan *et al*. 2018, Pradhan *et al*. 2019, Tripathi *et al*. 2020, Williams and Lorenz 2020). However, oestrogen did not affect the amount or exposure of the main cell wall carbohydrates (Fig S2A-E), and did not significantly impact on the section of pro-inflammatory cytokines (Fig S2F-I).

In addition to direct detection of cell wall PAMPs, *C. albicans* activates the alternative complement cascade resulting in the deposition of complement (C3) on its surface inducing opsonophagocytosis via the complement receptors CR1 and CR3. To investigate whether oestrogen-induce innate immune evasion was mediated via inhibition of complement, we assessed phagocytosis rates in heat-inactivated serum, where the major complement proteins (i.e. C3) are denatured. Overall, the phagocytosis rates of *C. albicans* in heat-inactivated serum were lower than in live serum, confirming the importance of opsonophagocytosis in the recognition of *C. albicans*. However, adaptation of *C. albicans* to oestrogen did not result in further evasion of phagocytosis in heat-inactivated serum (Fig 2A), suggesting that pre-exposure of *C. albicans* to oestrogen impacts complement activation. Supplementation of heat-inactivated serum with purified C3 restored oestrogen-induced immune evasion (Fig 2B), confirming that oestrogen-induced immune evasion likely occurs through the avoidance of opsonophagocytosis.

**Figure 2.**
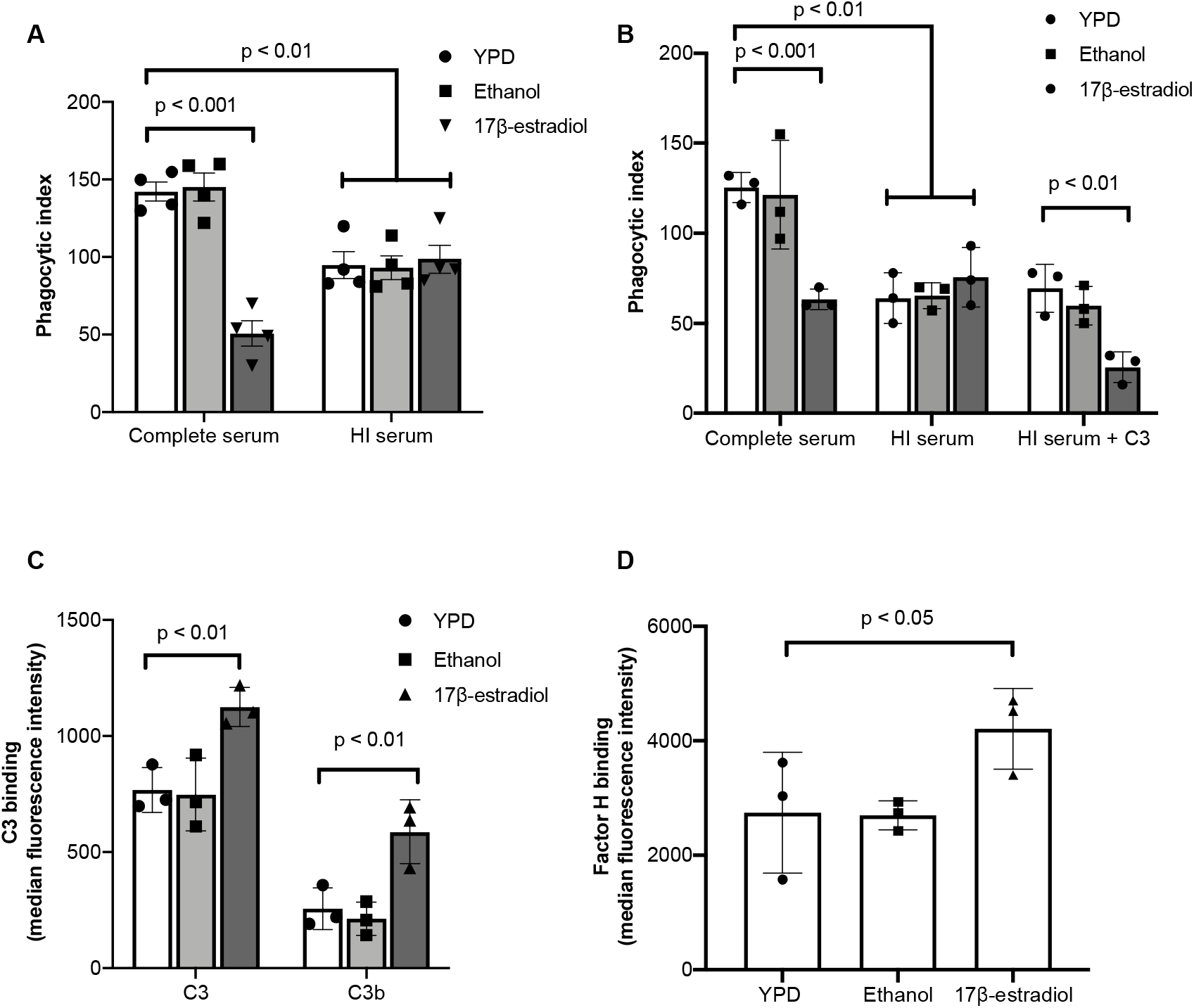
Oestrogen promotes innate immune evasion of *C. albicans* through the inhibition of opsonophagocytosis. **A)** J774A.1 macrophages were maintained in either complete serum or heat-inactivated (HI) serum and infected with *C. albicans* pre-exposed to YPD, 0.3% ethanol or 10 μM 17β-estradiol at a MOI of 5. **B)** J774A.1 macrophages were maintained in either complete serum, heat-inactivated (HI) serum, or heat-inactivated serum supplemented with purified C3 and infected with *C. albicans* pre-exposed to YPD, 0.3% ethanol or 10 μM 17β-estradiol at a MOI of 5. **C)** *C. albicans* was grown in YPD with or without 10 μM 17β-estradiol, incubated in human serum for 20 minutes, fixed with 4% PFA and C3 and C3b binding quantified by FACS. **(D)**. *C. albicans* was grown in YPD with or without 10 μM 17β-estradiol, incubated in human serum for 20 minutes, fixed with 4% PFA and Factor H binding quantified by FACS. Data represent the mean ± SEM from at least three independent experiments (also see Figure S2).

Next, we tested whether adaptation of *C. albicans* to oestrogen affected the deposition of C3 and C3b on the fungal cell surface. Oestrogen-adapted cells exhibited enhanced C3 and C3b deposition compared to non-adapted cells (Fig 2C), suggesting that adaptation to oestrogen promotes C3 and C3b binding on the *C. albicans* cell surface. Increased deposition of complement is normally associated with increased phagocytosis. However, processing of C3 can be inhibited by the recruitment of host regulatory proteins like Factor H, and several pathogens are known to avoid opsonophagocytosis through enhanced recruitment of Factor H (Dasari *et al*. 2018, van der Maten *et al*. 2018). Therefore, the ability of oestrogen adapted *C. albicans* cells to bind Factor H was quantified. Oestrogen-adapted cells bound significantly more Factor H, compared to the solvent control (Fig 2D), suggesting that oestrogen-adapted *C. albicans* evade the innate immune system through enhanced Factor H recruitment and decreased opsonophagocytosis.

### Bcr1 plays a key role in oestrogen-induced innate immune evasion

Altered deposition of complement on the fungal cell surface in response to oestrogen stimulation, suggests that oestrogen induces altered expression of *C. albicans* cell surface proteins. To elucidate how quickly *C. albicans* adapts to oestrogen, the fungus was grown in the presence of 10 μM 17β-estradiol for varying lengths of time before macrophage phagocytosis rates were quantified. This time course analysis confirmed that phagocytosis rates gradually reduced after 60 minutes of growth in the presence of oestrogen (Fig 3A), suggesting that *C. albicans* undergoes some form of transcriptional or translational response to oestrogen. To identify how oestrogen affects the global transcriptional profile of *C. albicans*, RNA-Seq was performed on *C. albicans* cells that had been adapted to 10 μM 17β-estradiol for 4 hours. Despite having a strong impact on the host-pathogen interaction, oestrogen only had a mild effect on the transcriptome of *C. albicans*, with 59 genes being moderately up-regulated and 75 genes being moderately down-regulated in the presence of oestrogen (Table S1, fold change > 1.5, FDR < 0.05). Genes that were upregulated were largely involved in hormone binding, RNA processing and FMN binding, while down regulated genes were largely involved in oxidoreductase activities (Table S2).

**Figure 3.**
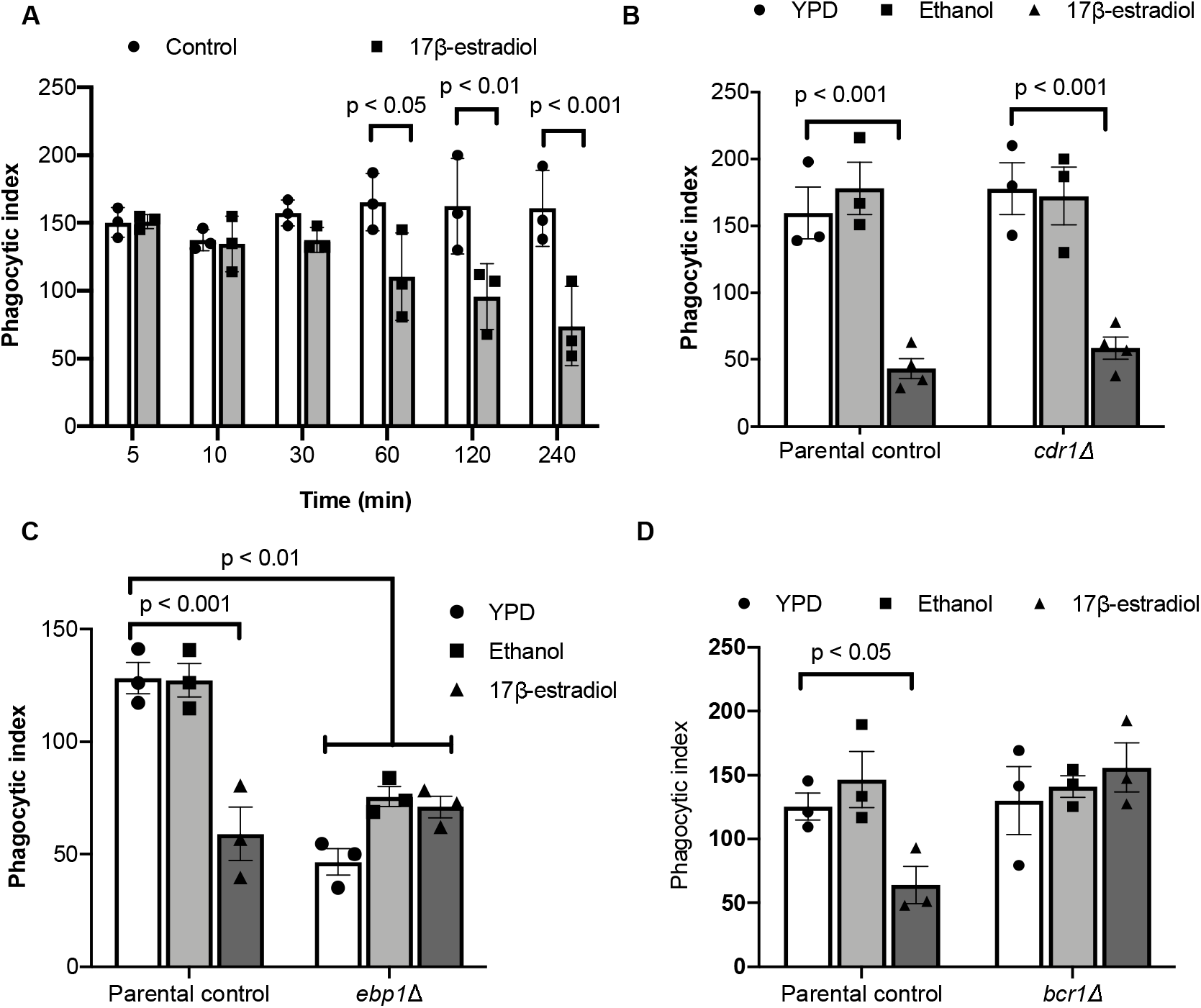
Oestrogen induces transcriptional changes in fungal cell wall proteins through Ebp1 and Bcr1. **A)** *C. albicans* was grown in YPD with or without 10 μM 17β-estradiol for increasing periods of time, cells were harvested, washed and co-incubated with J774A.1 macrophages for 45 minutes, and phagocytosis rates quantified. Data represent the mean and SEM for three independent biological experiments. **B)** The *cdr1*Δ. mutant and parental control strains were grown with or without 10 μM 17β-estradiol for 4 hrs and then exposed to J774A.1 macrophages for 45 minutes and phagocytosis rates quantified. **C)** The *ebp1*Δ mutant and parental control strains were grown with or without 10 μM 17β-estradiol for 4 hrs and then exposed to J774A.1 macrophages for 45 minutes and phagocytosis rates quantified. **D)** The *bcr1*Δ mutant and parental control strain were grown with or without 10 μM 17β-estradiol for 4 hrs and then exposed to J774A.1 macrophages for 45 minutes and phagocytosis rates quantified. All data represent the mean ± SEM from at least three independent biological experiments (also see Tables S1-5).

To identify whether the two differentially expressed genes involved in hormone binding (*EBP1* and *CDR1*) play a role in oestrogen induced innate immune evasion, the phagocytosis rates of the mutants in the presence and absence of oestrogen were quantified. Deletion of *CDR1* did not affect oestrogen induced innate immune evasion (Fig 3B). However, deletion of *EBP1* resulted in consistently low phagocytosis even in the absence of hormone stimulation (Fig 3C), suggesting that Ebp1 is a negative regulator of innate immune evasion.

To identify whether oestrogen mediates its affects through a conserved signalling pathway, the promoter regions of the differentially expressed genes were analysed for the presence of conserved transcription factor binding motifs. From the top three putative motifs identified in the 5’ UTR of the differentially expressed genes (Table S3), 18 documented transcription factor binding motifs were identified (Table S4). Given that phagocytosis is governed by interactions between PAMPs located on the fungal cell surface, and PRRs on the phagocyte, we reasoned that enhanced Factor H binding, and therefore, innate immune evasion is likely to be mediated by proteins in the cell wall. From the 18 identified transcription factors, only Bcr1 has been linked to regulation of the fungal cell surface (Nobile and Mitchell 2005). To identify whether Bcr1 plays a role in the oestrogen response, phagocytosis rates of the *bcr1*Δ mutant were quantified in the presence and absence of oestrogen. Deletion of *BCR1* resulted in the loss of innate immune evasion in the presence of oestrogen (Fig 3D), confirming that Bcr1 is critical for this process.

To identify putative target genes of Bcr1 we analysed gene expression data from Nobile and Mitchell (Nobile and Mitchell 2005) in combination with data available on Pathoyeastract (Monteiro *et al*. 2019). In total, 45 genes were identified with documented *BCR1* binding sites in their promoters. To identify Bcr1 targets that function in the cell wall, we performed GO term analysis on these 45 genes, the top three GO terms are “cell surface”, “plasma membrane” and fungal-type cell wall” (Table S5). Of these cell surface associated genes, only Gpd2 has been linked to interaction with the complement system (Luo *et al*. 2012).

### Oestrogen-induced innate immune evasion is mediated through Gpd2

*GPD2* is a glycerol-3-phosphate dehydrogenase, which has been shown to localise to the fungal cell surface in response to serum (Marín *et al*. 2015). Gpd2 has been classed as a moonlighting protein (a protein that has a biological function not predicted by its amino acid sequence) due to its ability to bind key regulatory components of the alternative complement system including Factor H, Factor H like protein-1 (FHL-1) and plasminogen (Luo *et al*. 2012). Thus, we hypothesised that oestrogen induced innate immune evasion of *C. albicans* is mediated via Gpd2 dependent acquisition of Factor H. To test this hypothesis, *GPD2* was deleted in *C. albicans* and phagocytosis rates in the presence and absence of oestrogen determined. Deletion of *GPD2* prevented oestrogen-dependent inhibition of macrophage phagocytosis rates (Fig 4A). To determine whether the loss of innate immune evasion in the *gpd2*Δ mutant correlated with a reduction of Factor H recruitment to the fungal cell surface, we quantified Factor H binding in the *gpd2*Δ mutant in the presence and absence of oestrogen. As predicted, the *gpd2*Δ mutant did not bind more Factor H than wild type cells in the presence of oestrogen (Fig 4B), confirming that immune evasion is dependent on Factor H recruitment. To confirm that enhanced expression of *GPD2* is sufficient to promote innate immune evasion, we over-expressed *GPD2* in *C. albicans*. Over-expression of *GPD2* (2.75-fold increased mRNA expression compared to wild type cells) resulted in reduced phagocytosis rates irrespective of oestrogen treatment (Fig 4C), thereby confirming that enhanced expression of *GPD2* is sufficient to promote *C. albicans* innate immune evasion. Given that deletion of *EBP1* results in constitutive innate immune evasion similar to the overexpression of *GPD2*, we quantified the expression of *GPD2* in the *ebp1*Δ mutant. Deletion of *EBP1* resulted in 2.74-fold higher expression of *GPD2* (comparable levels of *GPD2* expression to the over-expression strain) compared to wild type cells (Fig 4D), suggesting Ebp1 is a negative regulator of *GPD2* expression and therefore innate immune evasion.

**Figure 4.**
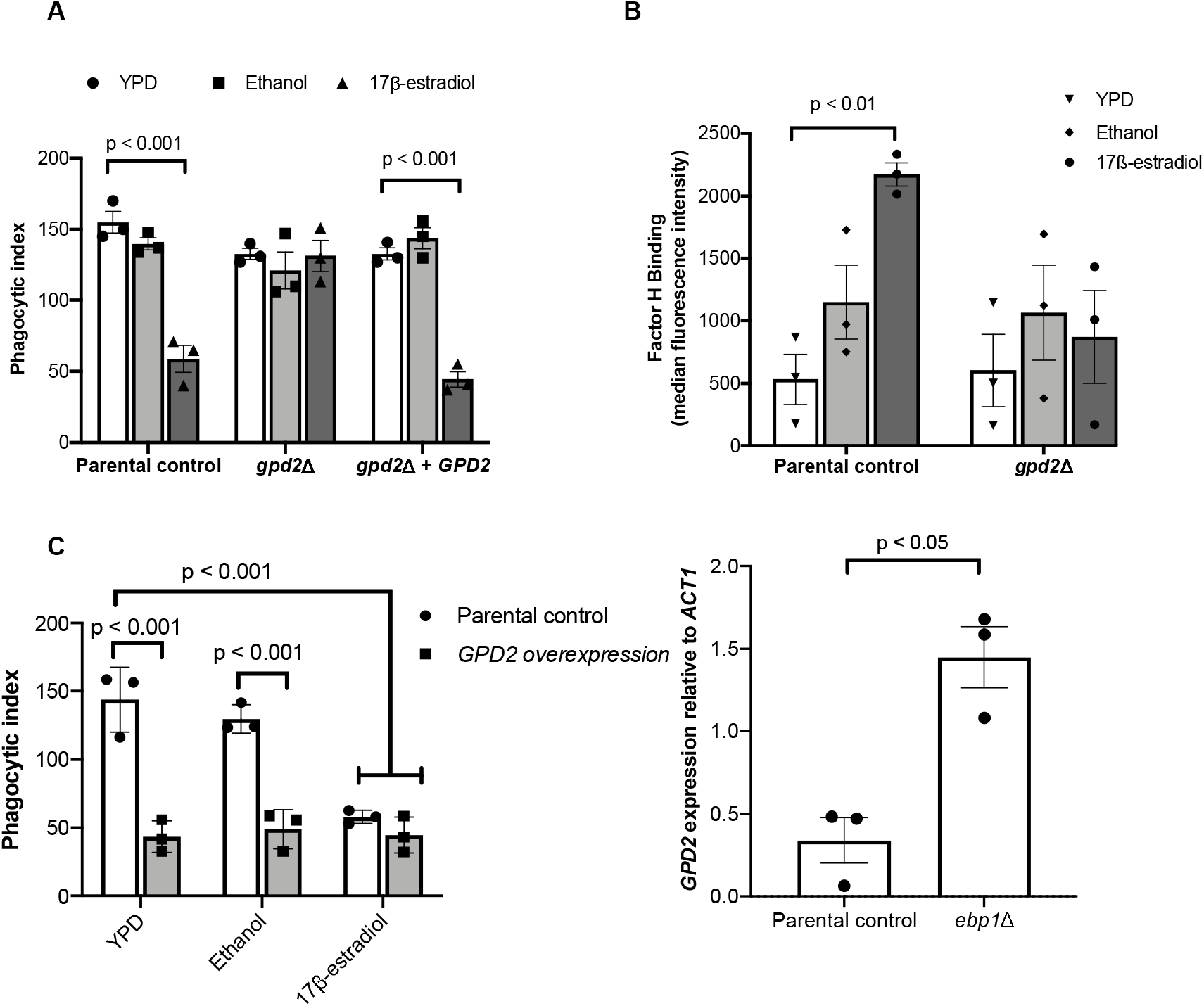
Inhibition of opsonophagocytosis is dependent on Gpd2. **A)** The parental control strain, *gpd2*Δ mutant and reconstituted control strains were grown in YPD with or without 10 μM 17β-estradiol for 4 hr, and then J77A.1 macrophage phagocytosis rates were quantified using ImageJ. **B)** *C. albicans* strains were grown in YPD with or without 10 μM 17β-estradiol, incubated in human serum for 20 minutes, fixed with PFA and Factor H binding quantified by FACS. **C)** The parental control (CAI4-pSM2) and *GPD2* overexpression (CAI4-pSM2-*GPD2*) strains were grown in YPD with or without 10 μM 17β-estradiol for 4 hr, washed and co-incubated with J774A.1 macrophages at an MOI = 5 for 45 min. Phagocytosis rates were quantified using ImageJ. **D)** The *ebp1*Δ mutant and the parental control strain were grown in YPD to mid-log phase, total RNA extracted and *GPD2* gene expression quantified by RT-PCR relative to *ACT1*. All data represent the mean ± SEM from at least three independent biological experiments.

### Oestrogen-induced immune evasion plays a key role in *C. albicans* pathogenicity

Having established that adaptation of *C. albicans* to oestrogen inhibits phagocytosis, we explored whether this phenomenon could influence virulence *in vivo* using a zebrafish larval model for disseminated disease. Previously, it was shown that exposing zebrafish larvae (3 hours post fertilisation) to media containing 1 μM oestrogen results in an *in vivo* oestrogen concentration of 0.0057 μM, equivalent to physiological levels during pregnancy in humans (Hao *et al*. 2013, Souder and Gorelick 2017). Taking advantage of this observation, zebrafish larvae were infected with *C. albicans* SC5314 and maintained in E3 media supplemented with 1 μM oestrogen and survival rate monitored for up to 5 days post fertilisation. Compared to zebrafish incubated in E3 media alone, or E3 media supplemented with ethanol, supplementation of E3 media with 1 μM oestrogen enhanced the virulence of *C. albicans*, resulting in an 63% reduction in zebrafish survival 60 hrs post infection (Fig 5A).

**Figure 5.**
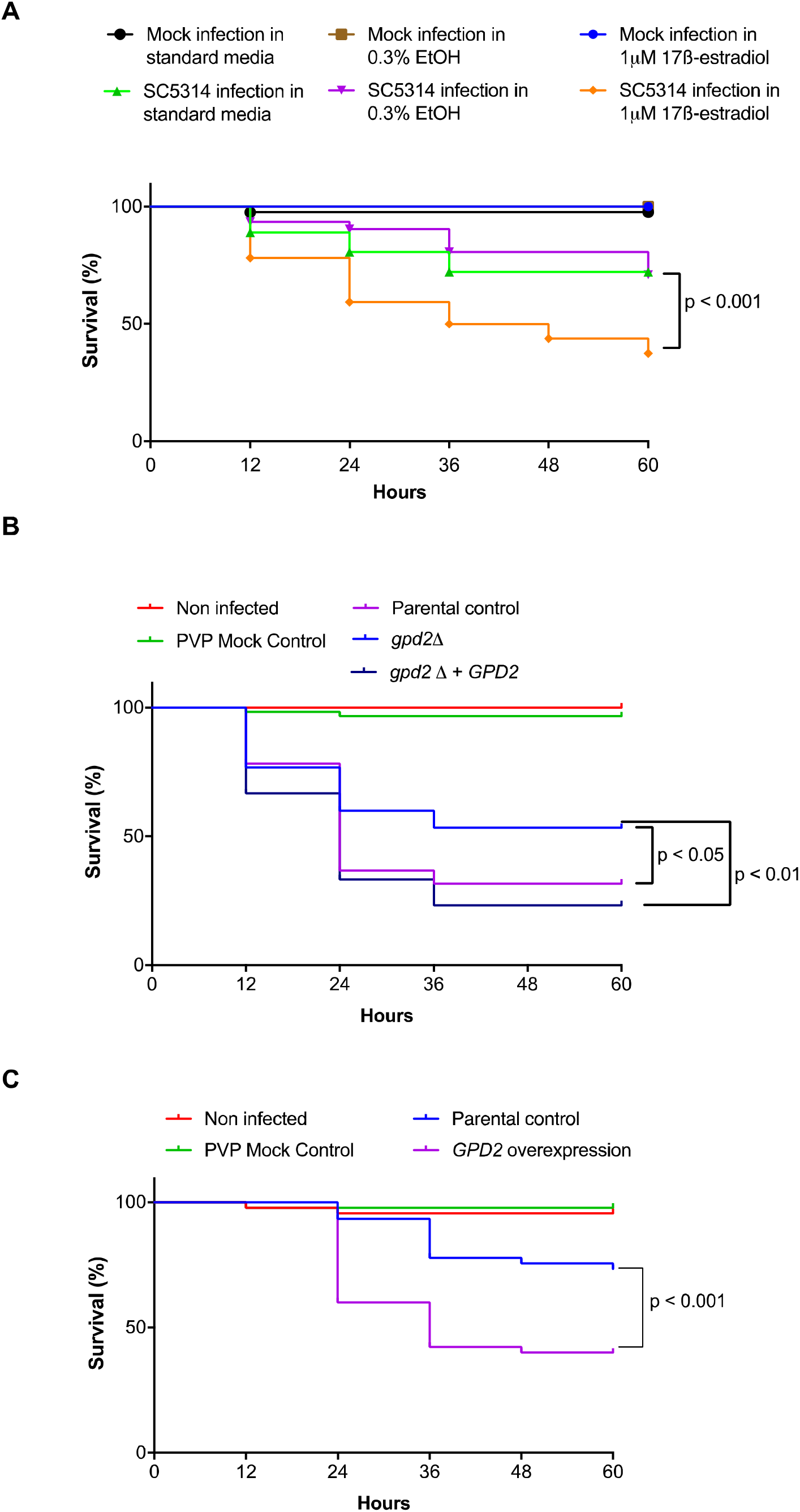
Oestrogen promotes the virulence of *C. albicans* in a zebrafish larval infection model. **A)** *C. albicans* (SC5314) were microinjected into the hindbrain ventricle of zebrafish larvae in the Prim25 stage. Infected larvae were maintained in E3 media, or E3 media supplemented with 0.3% ethanol, or 1 μM 17β-estradiol and larval survival monitored every 24 h until day 5 post fertilisation. **B**) The parental control (SN250-CIP30), *gpd2*Δ mutant (*gpd2*Δ-CIP30) and reconstituted control (*gpd2*Δ-CIP30-*GPD2*) strains were microinjected into the hindbrain ventricle of zebrafish larvae in the Prim25 stage dpf. Larvae were maintained in E3 media and larval survival monitored every 24 h until day 5 post fertilisation. **C)** The parental control (CAI4-pSM2) or GDP2 overexpression (CAI4-pSM2-*GPD2*) strains were microinjected into the hindbrain ventricle of zebrafish larvae in the Prim25 stage dpf. Larvae were maintained in E3 media and larval survival monitored every 24 h until day 5 post fertilisation. The survival curves represent data pooled from three independent experiments. Statistically significant differences were determined by Log-rank (Mantel-Cox) test.

To investigate whether Gpd2 is required for virulence in this model, zebrafish were infected with the parental control strain, the *gpd2*Δ mutant, or reconstituted control strains of *C. albicans*. Deletion of *GPD2* led to attenuation of *C. albicans* virulence, which was restored to parental control levels via complementation with a single copy of *GPD2* (Fig 5B). To assess whether increased expression of Gpd2 is sufficient to drive fungal virulence, *GPD2* was over expressed in the CAI4 background. Infection of zebrafish larvae with *C. albicans* over-expressing *GPD2* enhanced *C. albicans* virulence in the absence of oestrogen compared to the respective control strain (Fig 5C). Therefore, Gpd2 plays a key role in promoting *C. albicans* virulence *in vivo*.

## Discussion

Vulvovaginal candidiasis (VVC) is a mucosal infection affecting 75% of the female population of reproductive age (SOBEL 1988). Oestrogen is known to govern susceptibility to VVC infections, with women with low circulatory oestrogen levels (i.e. postmenopausal women) having a low risk of developing VVC, and women with high oestrogen levels (i.e. during pregnancy, or women taking high oestrogen containing oral contracepts) having a high risk of VVC. Elevated oestrogen levels increase glycogen at the vaginal mucosa, reduce leukocyte infiltration, and reduce antifungal activity of epithelial cells, promoting infection (Fidel *et al*. 2000, Dennerstein and Ellis 2001, Salinas-Muñoz *et al*. 2018). However, here we show that, in addition to these effects on the host, oestrogen promotes adaptation responses in *C. albicans* which induce evasion of the innate immune system through Gpd2-dependent inhibition of complement mediated opsonophagocytosis. Unlike the induction of fungal morphogenesis, which appears to be limited to 17β-estradiol (Cheng *et al*. 2006), all tested forms of oestrogen promoted innate immune evasion, suggesting that *C. albicans* has evolved at least two signalling pathways that are responsive to oestrogen.

Complement evasion is a successful mechanism employed by viruses, bacteria, parasites and fungi to escape the innate immune system (Lambris *et al*. 2008). Evasion of the complement system can be mediated through the secretion of degradative proteins, or by recruitment of host regulatory proteins (Merle *et al*. 2015). One of the most common methods pathogens use to evade complement is through enhanced recruitment of Factor H, a key regulatory protein in the complement system, to the microbial surface. Bound Factor H prevents further activation of the alternative complement system through both the destabilisation of the C3 convertase and enhancement of Factor I mediated degradation of C3b to iC3b, reducing opsonisation and inhibiting the formation of the membrane attack complex (Merle *et al*. 2015). *C. albicans* has been shown by others to recruit Factor H to its surface through the expression of moonlighting proteins. Moonlighting proteins are proteins that perform a biological function not predicted from the amino acid sequence. So far, four moonlighting proteins (Phr1, Gpd2, Hgt1 and Gpm1) have been identified in *C. albicans* (Poltermann *et al*. 2007, Luo *et al*. 2010, Luo *et al*. 2012, Kenno *et al*. 2018). Although Gpd2 is predicted to be a cytoplasmic protein involved in cellular redox, Gpd2 has also been identified in cell wall proteomic studies (Marín *et al*. 2015), suggesting that Gpd2 functions in the cell wall. Interestingly, Gpd2 was only identified in the cell wall proteome of *C. albicans* grown in complete serum, and not heat-inactivated serum where complement proteins have been inactivated (Marín *et al*. 2015). Therefore, deposition of complement on the fungal cell surface may drive localisation of Gpd2 to the cell surface.

Purified Gpd2 binds all three complement regulatory proteins (Luo *et al*. 2012), although the biological significance of these interactions in infections is not known. Factor H and FHL1 bound to Gpd2 remain active, cleaving C3b and thereby inhibiting the alternative complement system (Luo *et al*. 2012). In addition, Gpd2 can also bind plasminogen, which is then processed into plasmin, contributing to inactivation of the alternative complement system (Luo *et al*. 2012). FACS analysis of the deposition of complement proteins on the surface of *C. albicans* in response to oestrogen suggested that oestrogen-adapted cells bound more Factor H, indicative of oestrogen promoting complement evasion. However, oestrogen-adapted *C. albicans* also bound more C3 and C3b, which is surprising as binding of Factor H should prevent C3 deposition. Factor H is a glycoprotein that is made up of 20 complement control proteins (CCP). Microbes bind Factor H at various positions, with *Neisseria* species binding Factor H at CCP6-8 (Ngampasutadol *et al*. 2008). However, multiple microbes bind Factor H at CCP20, and this has been termed the Common Microbial Binding site (Meri *et al*. 2013). Binding of Factor H to some of these microbial cell surface proteins at CCP20 results in increased affinity of Factor H for C3b (Meri *et al*. 2013). The enhanced affinity of Factor H for C3b might explain why we observe elevated binding of both C3b and Factor H, and would suggest that Gpd2 binds to CCP20. The formation of this stable tri-partite complex (microbial protein, Factor H, C3b) results in enhanced activity of Factor H and therefore rapid inactivation of the alternative complement cascade (Meri *et al*. 2013), which would explain why we observe reduced levels of phagocytosis (Fig 6).

**Figure 6.**
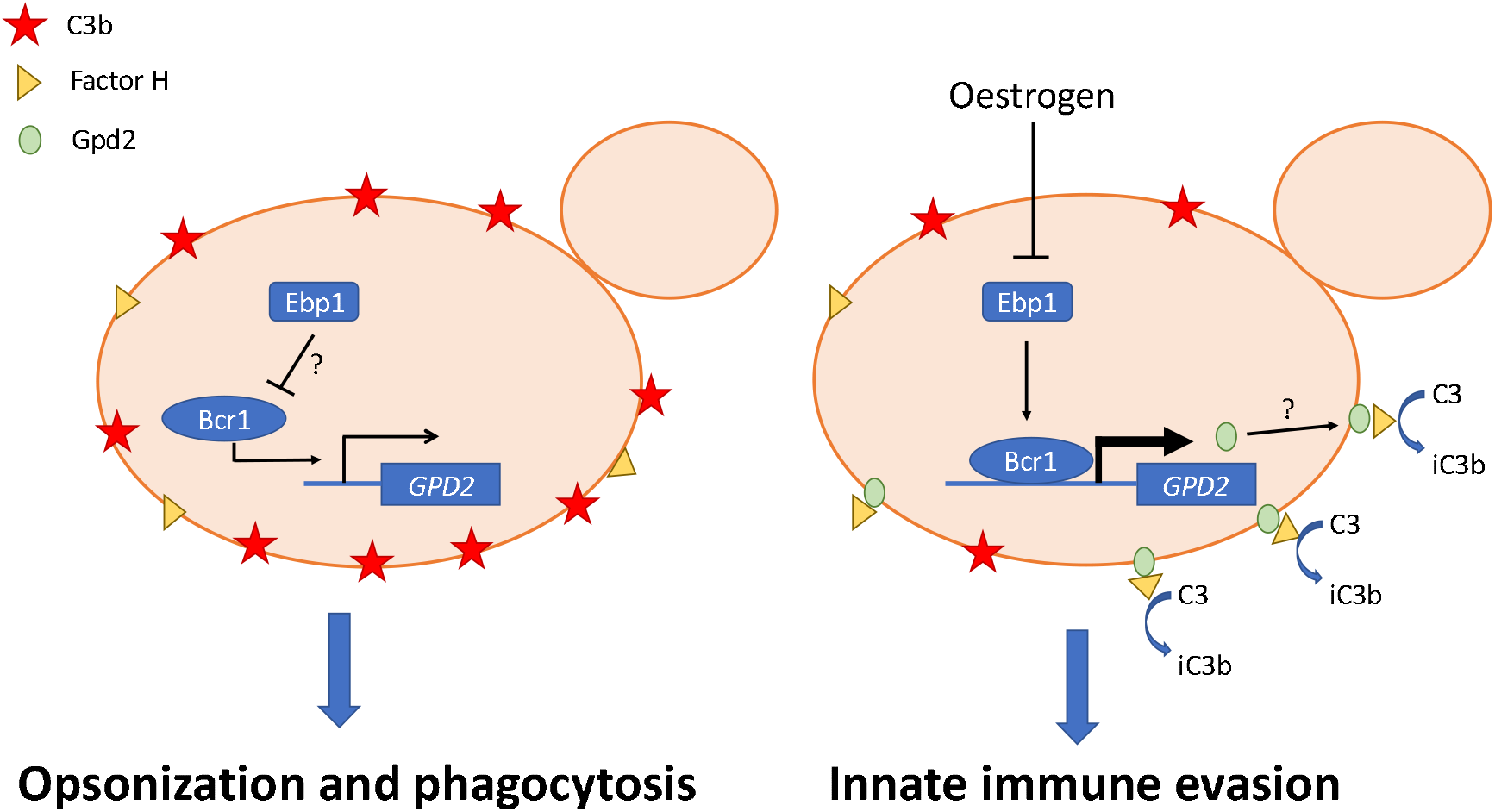
Proposed mechanism of oestrogen-induced innate immune evasion. Under standard laboratory conditions the C3 is deposited on the surface of *C. albicans* resulting in effective phagocytosis. However, in the presence of oestrogen the NADPH activity of Ebp1 is reduced resulting in derepression (through an as yet to be identified signalling pathway) and surface localisation of Gpd2. Gpd2 recruits Factor H to the fungal cell surface, preventing the formation of the C3 convertase and disposition of C3 on the fungal cell surface. Reduced opsonisation results in innate immune evasion and insufficient clearance of *C. albicans*, promoting fungal pathogenicity.

Innate immune evasion was only observed after *C. albicans* had been grown in the presence of oestrogen for 60 minutes, suggesting that the response required either transcriptional or post-transcriptional regulation. However, in line with previous studies (Cheng *et al*. 2006), global transcriptional analysis confirmed that only a small proportion of *C. albicans* genes were differentially regulated by oestrogen. In comparison to the work of Cheng *et al*., where most of the differentially regulated genes are involved in fungal morphogenesis (Cheng *et al*. 2006), our analysis mainly identified genes involved in metabolism and oxidation-reduction processes. This lack of overlap is likely reflective of the growth conditions used, as Cheng *et al*., used RPMI where oestrogen promotes hyphal formation (Cheng *et al*. 2006, Salinas-Muñoz *et al*. 2018), while under our experimental conditions (YPD) oestrogen did not promote hyphal formation. Although innate immune evasion was dependent on the presence of Gpd2, *GPD2* mRNA levels were only moderately increased in response to oestrogen, suggesting that other factors (i.e. post-translational modifications, protein localisation) may be involved. Given that Gpd2 has only been identified as a cell wall protein under specific conditions (Marín *et al*. 2015), it might be that in response to oestrogen a higher level of Gpd2 is recruited to the cell wall, enabling the fungus to bind more Factor H. In *S. cerevisiae*, the activity of both Gpd1 and Gpd2 is regulated via post-translational modification (Lee *et al*. 2012). Snf1-dependent phosphorylation of *Sc*Gpd1 (orthologue of *CaGPD2*) results in decreased enzyme activity (Lee *et al*. 2012). Although *Ca*Gpd2 has several predicted phosphorylation sites, these are located in different regions of the protein compared to *Sc*Gpd1, and therefore it is possible that phosphorylation/dephosphorylation at any of these sites in *Ca*Gpd2 could affect the localisation, or potential to bind Factor H.

Ebp1, despite showing no homology to mammalian hormone binding proteins, has been described as an oestrogen binding protein. Ebp1 is homologous to old yellow enzyme 2 (*OYE2*) of *S. cerevisiae*, an NADPH oxidoreductase. Biochemical characterisation of recombinant Ebp1 confirms that Ebp1 is an NADPH oxidoreductase, and that oestrogen inhibits this enzymatic activity (Madani *et al*. 1994, Buckman and Miller 1998). The importance of Ebp1 enzymatic activity in *C. albicans* pathogenicity is not known, but here we demonstrate that Ebp1 may serve as a negative regulator of innate immune evasion through the regulation of *GPD2*. Interaction of oestrogen with Ebp1, through an as yet unidentified pathway likely involving Bcr1, results in elevated expression of *GPD2* and, potentially in association with other post-translational modifications, promotes innate immune evasion and pathogenicity (Fig 6).

In recent years zebrafish larvae have become an excellent model for studying pathogenicity mechanisms. Zebrafish have a complement system that is structurally and functionally similar to the mammalian complement system (Zhang and Cui 2014), and an orthologue of Factor H has been identified and cloned (Sun *et al*. 2010). The presence of oestrogen in the zebrafish larval model of disseminated *C. albicans* infection, confirmed that oestrogen promotes the virulence of *C. albicans*, suggesting that oestrogen dependent inactivation of the alternative complement system promotes the virulence of *C. albicans in vivo*. In support of this, deletion of *GPD2* attenuated fungal virulence, while over expression of *GPD2* enhanced virulence in the zebrafish model of infection, confirming that Gpd2 is an important fungal virulence factor. Given the strong association of elevated oestrogen levels with the development of VVC, it is important to understand the role of Gpd2 in promoting innate immune evasion, and may lead to the development of alternative treatment options for VVC and RVVC.

## Acknowledgements

We would like to thank the fungal community for the donation of fungal strains, Prof G. Brown (University of Exeter) for FC-Dectin-1, Prof G. Ramage for clinical isolates, the Environmental Omics facility at the University of Birmingham for help with the RNA sequencing, and Elizabeth Ballou (University of Birmingham) for critical reading of the manuscript, and Prof A. Mitchell for providing access to raw data from (Nobile and Mitchell 2005). PK is funded by the Wellcome trust strategic award for medical mycology (097377/Z/11/Z), FC and RAH are funded by the MRC (MR/L00903X/1) and BBSRC (BB/R00966X/1).

## Author Contributions

PK, FC, BK, HG, conducted the experiments, FC, HT conducted the bioinformatic analysis, RAH conceived the idea and designed the experiments, RAH and HT wrote the paper.

## STAR Methods

## Resource availability

### Lead Contact

Further information and requests for resources and reagents should be directed to and will be fulfilled by the lead contact Dr R Hall.

### Materials Availability

New strains and materials made during this study will be made available upon request to the lead contact Dr R Hall subject to the completion of a materials transfer agreement.

### Data and Code Availability

The raw RNA Seq data files are available at the Gene Expression Omnibus (GEO) database (http://www.ncbi.nlm.nih.gov/geo/) at the following accession number GSE145240.

## Experimental Model and Subject Details

### Fungi

*Candida* strains (Table S6) were routinely maintained on YPD agar (1% yeast extract, 1% peptone, 2% glucose and 2% agar). For broth cultures all strains were cultured in YPD (1% yeast extract, 1% bacto-peptone, 2% glucose) buffered to pH6 with 3.57% HEPES. Oestrogen was diluted in 10% ethanol to a stock concentration of 100 μg/ml and diluted into YPD at the required concentrations, maintaining the final ethanol concentration at 0.3%.

### Cell Lines

J774A.1 macrophages (Sigma, UK) were maintained in low glucose DMEM supplemented with 5% FBS, 2mM L-glutamine and 2 mM Pen/Strep, 37C, 5% CO_2_.

### Zebrafish

Zebrafish care and experiment protocols were performed in accordance with the Home Office project license 40/3681 and personal license IE905E215 as per Animal Scientific Procedures Act 1986. Wild type (AB) *Danio rerio* zebrafish used in the study were housed in a recirculating system of gallon tanks at the University of Birmingham Zebrafish Facility. To obtain embryos, 4 male and 5 female fish were transferred into a breeding tank and maintained at 28°C, 14 h light/10 h dark cycle. Embryos were collected the following day, sorted and incubated at 30°C for 24 h in E3 media (5 mM NaCl, 0.17 mM KCl, 0.33 mM CaCl_2_, 0.33 mM MgSO_4_, 0.00003% methylene blue, pH 7). Embryos were maintained at a density of 100 per 14.5 cm dish containing 150 mL E3 media supplemented with 0.02 mg/mL Phenylthiourea.

### Human macrophages and monocytes

Protocols for human blood collection and isolation of neutrophils and peripheral blood mononuclear cells (PBMCs) were approved by ethical review board of the School of Biosciences at the University of Birmingham. Blood was collected anonymously and on voluntary basis after getting written informed consent.

## Method Details

### Phagocytosis experiments

Phagocytosis assay was performed as previously described (Cottier *et al*. 2019). Briefly, overnight cultures of *C. albicans* were sub-cultured 1:100 in fresh YPD media, media supplemented with 0.3% ethanol, or media supplemented with oestrogen (0.0001 μM, 0.01 μM or 10 μM) and incubated at 37 C, 200 rpm for 4 h. *C. albicans* cells were washed three times in PBS and 1 x10^5^ J774A.1 macrophages (Sigma, UK) were infected with 5 x 10^5^ yeast cells (multiplicity of infection [MOI] of 5) for 45 min at 37°C, 5% CO_2_. Cells were washed with PBS to remove non-phagocytosed yeast cells, and phagocytosis stopped by fixing with 4% paraformaldehyde (PFA) for 20 minutes. Samples were stained for 30 minutes with 50 μg/ml ConA-FITC to stain non-phagocytosed fungal cells, washed and imaged. Phagocytosis events were scored from multiple fields of view using imageJ. When required, J774A.1 cells were maintained in DMEM supplemented with heat inactivated serum and the assay was complemented with 1 μg/mL C3 (Sigma, C2910) or C3b (Sigma, 204860).

### Human macrophages and neutrophils

PBMCs and neutrophils were isolated as previously described (Sherrington *et al*. 2017). Neutrophils were seeded at 2 x 10^5^ cells/mL in 24-well plates in serum free RPMI supplemented with 100 mM L-glutamine, incubated for 1 h at 37°C, 5% CO_2_ and then co-incubated with 1 x 10^6^ *C. albicans* cells (MOI = 5) for 45 min at 37°C, 5% CO_2_. Cells were immediately fixed with 4% PFA and stained with 50 μg/ml ConA-FITC for 30 minutes to stain non-phagocytosed fungal cells, washed and imaged. Phagocytosis events were scored from multiple fields of view using imageJ. To assess primary macrophage phagocytosis rates PBMCs (0.5 x10^6^) were seeded into 24 well plates in differentiation media (RPMI 1640 supplemented with 100 mM L-glutamine, 10% human AB serum, 100 mM Pen/Strep and 20 ng/ml M-CSF) for 7 days replacing the media every 2-3 days, and then phagocytosis rates determined as described above. To assess cytokine production 2.5 x10^4^ PBMCs were stimulated with 5 x10^4^ PFA fixed *C. albicans* for 24 hours, supernatants collected and stored at −20°C and cytokine concentrations quantified by ELISA.

### Genetic manipulation of C. albicans

All primers used in genetic manipulation of *C. albicans* are listed in S2 Table. To reintroduce *GPD2* into *C. albicans gpd2*Δ mutant, the *GPD2* locus was PCR-amplified from *C. albicans* SC5314 genomic DNA using primers GPD2-SacI-F and GPD2-NotI-R. The PCR product was cloned into CIp30 plasmid (Dennison *et al*. 2005) restricted with *SacI* and *NotI* using T4 DNA ligase. The generated CIp30-*GPD2* was linearized with *StuI* and transformed into *C. albicans gpd2*Δ mutant by standard heat-shock transformation (Walther and Wendland 2003).

To generate a *C. albicans* strain that over expresses *GPD2*, the open reading frame of *GPD2* was cloned into pSM2 (Barkani *et al*. 2000) using primers GPD2-OE-F and GPD2-OE-R under the control of the *TEF2* promoter. The resulting plasmid was linearized with *PacI*, and integrated into CAI4 at the *URA3* locus by standard heat-shock transformation.

To generate *ebp1*Δ, 500 bp of the 5’ and 3’ UTR were amplified from genomic DNA using primers EBP1-5F, EBP1-5R, and EBP1-3F and EBP1-3R. The resulting PCR products were purified, digested with *EcoRV* and *SacI* or *HindIII* with *KpnI* and cloned into the mini URA blaster cassette pDDB57 (Wilson *et al*. 2000). The resulting disruption cassette was digested with *EcoRV* and *KpnI* and transformed in CAI4 by standard heat-shock transformation. Resulting colonies were screened by PCR and positive colonies were plated onto YNB supplemented with 5-fluoroorotic acid (5-FOA) and uridine to select for spontaneous homologous recombination and loss of *URA3*. Resulting colonies were then re-transformed with the *EBP1* knockout cassette, and loss of both alleles was confirmed by PCR, and lack of expression was confirmed by qPCR. The URA baster cassette was recycled and *URA3* replaced at its native locus by transforming the strain with pSM2.

### *Immunofluorescent staining of* C. albicans *cell wall components*

*C. albicans* cells were stained as previously described (Sherrington *et al*. 2017). Briefly, *C. albicans* cells from overnight culture were sub-cultured in YPD broth with or without oestrogen supplementation and incubated at 37°C, 200 rpm for 4 h. Cells were harvested by centrifugation, washed in PBS and fixed with 4% PFA. To quantify total mannan, glucan and chitin levels in the cell wall, cells were stained with 50 μg/ml TRITC-conjugated concanavalin A (Molecular Probes, Life Technologies), 33.3 μg/ml Aniline Blue fluorochrome (Bioscience supplies) and 3 μg/ml Calcofluor White for 30 minutes. To quantify surface exposure of β1,3-glucan and chitin, cells were stained with 3 μg/ml Fc-Dectin-1 (a gift from G. Brown, University of Aberdeen) and 50 μg/ml TRITC-conjugated wheat germ agglutinin (Molecular Probes, Life Technologies).

### RNA sequencing

*C. albicans* cells were grown for 4 h in YPD broth with or without 10 μM 17β-estradiol at 37 C, 200 rpm. Cells were harvested by centrifugation, washed three times in PBS, and snapped frozen in liquid nitrogen. Total RNA was extracted as per manufacturer’s instructions using the RNeasy Plus mini kit (Qiagen). Total RNA was quantified using NanoDrop 8000 spectrophotometer (ND-8000-GL; Thermo Fisher). RNA samples were assessed for genomic DNA contamination by PCR and agarose gel electrophoresis. Samples were then processed as previously reported (Cottier *et al*. 2019). Sequencing reads are available at the Gene Expression Omnibus (GEO) database (http://www.ncbi.nlm.nih.gov/geo/) at the following accession number GSE145240.

Reads were analysed following a previous published method (Cottier *et al*. 2019) and using CLC Genomic workbench 11.0.1 software (Qiagen). In summary, adapter and quality trimming was performed before reads were mapped to *C. albicans* reference genome (Assembly 21, version s02-m09-r10). Transcript Per Kilobase Million (TPM) were reported for each open reading frame (ORFs). Statistical analysis was performed after addition to all values of the lowest TPM measurement, then data were log10 transformed and differential expression between condition were considered significant if the absolute value Fold Change >1.5 and FDR < 0.05. Pathoyeastract GO term finder (Inglis *et al*. 2012) was used to perform Gene ontology (GO) analysis with P-values corresponding to Bonferroni-corrected hypergeometric test P-values. Motif analyses were performed in MEME suite website (http://meme-suite.org/).

### RT-PCR

*C. albicans* cells were grown for 4 h in YPD broth with or without 10 μM 17β-estradiol at 37 C, 200 rpm. Cells were harvested by centrifugation, and snapped frozen in liquid nitrogen. RNA was extracted using the RNeasy Plus mini kit (Qiagen) as per manufacturer’s instructions. RNA quality and quantity were checked by electrophoresis and spectroscopy. The qRT-PCR was performed using the 2x qPCRBIO SyGreen mix kit (PCRbiosystem) according to manufacturer’s recommendations with 50 ng of total RNA (primers shown in Table S7). Relative quantification of gene expression was determined by the Delta Delta Ct method with *ACT1* as an endogenous control. mRNA expression was performed in technical triplicate, and data represent the mean and SEM from three independent biological repeats, and were analysed using a paired T-test with 95% confidence.

### Complement binding

*C. albicans* cells were grown for 4 h in YPD broth with or without 10 μM 17β-estradiol at 37 C, 200 rpm. Cells were harvested, washed three times in PBS, fixed on ice for 30 min with 4% PFA. About 2 x 10^6^ yeast cells were labelled with 400 μL 10% normal human serum for 20 min at 37 C, 200 rpm. Cells were washed thrice in PBS and incubated on ice with 100 μL of either 10 μg/mL Anti-Factor H Goat pAb (Sigma, 341276-1ML) or 1 μg/mL Goat anti Chicken IgY (H+L) diluted in 1% BSA/PBS. Cells were washed thrice in PBS and incubated in dark with 100 μL of either Rabbit anti Goat IgG (H+L) Secondary Antibody, Alexa Fluor 488 (Invitrogen, A11078) or Goat anti Chicken IgY (H+L) Secondary Antibody, Alexa Fluor 594, (Invitrogen, A11042) diluted 1:200 in 1% BSA/PBS. Cells were analysed by flow cytometry and median fluorescence intensity determined. FACS data were analysed by Kruskal-Wallis test followed by a post-hoc Dunn’s multiple comparisons test at 95% confidence.

### Zebrafish infection

Hind brain infections were performed as previously described (Mallick *et al*. 2016). Briefly, zebrafish at the prim25 stage were manually dechorionated, and anesthetized in 160 μg/mL Tricaine. Approximately 5 nL of injection buffer (10% PVP-40 in PBS, 0.05% phenol red) or *C. albicans* suspension at 5 × 10^7^ cells/mL in injection buffer was microinjected into the hindbrain ventricle via the otic vesicle to achieve a dose of 20-50 yeast/larva. Within 1 h of infection, larvae were screened by microscopy to remove fish noticeably traumatised from microinjection and to ascertain correct injection site and inoculum size. At least 15 larvae per condition were transferred into a 6-well plate, incubated at 28°C in E3 media either with or without 1 μM oestrogen and observed for survival every 24 h until day 5 post fertilisation.

## Quantification of Statistical Analysis

Unless indicated otherwise, data were analysed in Prism (version 8) and data presented in graphs represent the mean +/- SEM from at least three independent biological experiments, and the individual biological replicates are displayed on each graph. For phagocytosis data all experiments were performed in technical duplicates, with a minimum of three independent biological repeats. For each technical repeat multiple fields of view were imaged and the mean phagocytosis rate quantified. The mean from each independent biological replicate was then analysed by Kruskal-Wallis test followed by a post-hoc Dunn’s multiple comparisons test at 95% confidence. For quantification of immunofluorescence 10,000 cells were analysed by flow cytometry and median fluorescence intensity (MFI) quantified. FACS data were analysed by Kruskal-Wallis test followed by a post-hoc Dunn’s multiple comparisons test at 95% confidence. For zebrafish infections, data from three independent biological replicates were pooled together to determine percent survival. Data were analysed by Log-rank Mantel-Cox test and Gehan-Breslow-Wilcoxon test (extra weight for early time points).

## Supplemental Information

**Table S1. Differentially regulated genes identified by RNA Seq** (Excel file)

## References

Amirshahi, A., C. Wan, K. Beagley, J. Latter, I. Symonds and P. Timms (2011). “Modulation of the *Chlamydia trachomatis* in vitro transcriptome response by the sex hormones estradiol and progesterone.” BMC Microbiol 11: 150.

Ballou, E. R., G. M. Avelar, D. S. Childers, J. Mackie, J. M. Bain, J. Wagener, S. L. Kastora, M. D. Panea, S. E. Hardison, L. A. Walker, et al. (2016). “Lactate signalling regulates fungal ß-glucan masking and immune evasion.” Nature Microbiology 2: 16238.

Barkani, A. E., O. Kurzai, W. A. Fonzi, R. A. Porta, M. Frosch and F. A. Muhlschlegel (2000). “Dominant Active Alleles of *RIM101 (PRR2)* Bypass the pH Restriction on Filamentation of *Candida albicans*.” Mol. and Cell. Biol. 20: 4635–4647.

Buckman, J. and S. M. Miller (1998). “Binding and reactivity of *Candida albicans* estrogen binding protein with steroid and other substrates.” Biochemistry 37: 14326–14336.

Cheng, G., K. M. Yeater and L. L. Hoyer (2006). “Cellular and molecular biology of *Candida albicans* estrogen response.” Eukaryotic cell 5: 180–191.

Chotirmall, S. H., S. G. Smith, C. Gunaratnam, S. Cosgrove, B. D. Dimitrov, S. J. O’Neill, B. J. Harvey, C. M. Greene and N. G. McElvaney (2012). “Effect of estrogen on *Pseudomonas* mucoidy and exacerbations in cystic fibrosis.” N Engl J Med 366: 1978–1986.

Cottier, F., S. Sherrington, S. Cockerill, V. del Olmo Toledo, S. Kissane, H. Tournu, L. Orsini, G. E. Palmer, J. C. Pérez and R. A. Hall (2019). “Remasking of Candida albicans ß-Glucan in Response to Environmental pH Is Regulated by Quorum Sensing.” mBio 10: e02347–02319.

Dasari, P., I. A. Shopova, M. Stroe, D. Wartenberg, H. Martin-Dahse, N. Beyersdorf, P. Hortschansky, S. Dietrich, Z. Cseresnyés, M. T. Figge, et al. (2018). “Aspf2 From *Aspergillus fumigatus* Recruits Human Immune Regulators for Immune Evasion and Cell Damage.” Frontiers in immunology 9: 1635–1635.

Dennerstein, G. J. and D. H. Ellis (2001). “Oestrogen, glycogen and vaginal candidiasis.” Australian and New Zealand Journal of Obstetrics and Gynaecology 41: 326–328.

Dennison, P. M. J., M. Ramsdale, C. L. Manson and A. J. P. Brown (2005). “Gene disruption in *Candida albicans* using a synthetic, codon-optimised Cre-loxP system.” Fungal Genetics and Biology 42: 737–748.

Fidel, P. L., Jr., J. Cutright and C. Steele (2000). “Effects of reproductive hormones on experimental vaginal candidiasis.” Infection and immunity 68: 651–657.

Fonzi, W. A. and M. Y. Irwin (1993). “Isogenic Strain Construction and Gene Mapping in *Candida albicans*.” Genetics 134: 717–728.

García-Gómez, E., B. González-Pedrajo and I. Camacho-Arroyo (2013). “Role of sex steroid hormones in bacterial-host interactions.” Biomed Res Int 2013: 928290.

Ghazeeri, G., L. Abdullah and O. Abbas (2011). “Immunological differences in women compared with men: overview and contributing factors.” Am J Reprod Immunol 66: 163–169.

Gillum, A. M., E. Y. H. Tsay and D. R. Kirsch (1984). “Isolation of the *Candida albicans* gene for orotidine-5’-phosphate decarboxylase by complementation of *S. cerevisiae ura3* and *E. coli* pyrF mutations.” Mol. and Gen. Genet. 198: 179–182.

Gonçalves, B., C. Ferreira, C. T. Alves, M. Henriques, J. Azeredo and S. Silva (2016). “Vulvovaginal candidiasis: Epidemiology, microbiology and risk factors.” Critical reviews in microbiology 42: 905–927.

György, C. (2017). “Is there a hormonal regulation of phagocytosis at unicellular and multicellular levels? A critical review.” Acta Microbiologica et Immunologica Hungarica 64: 357–372.

Hall, R. A. (2015). “Dressed to impress: impact of environmental adaptation on the *Candida albicans* cell wall.” Molecular Microbiology 97: 7–17.

Hall, R. A., L. De Sordi, D. M. MacCallum, H. Topal, R. Eaton, J. W. Bloor, G. K. Robinson, L. R. Levin, J. Buck, Y. Wang, et al. (2010). “CO2 acts as a signalling molecule in populations of the fungal pathogen *Candida albicans*.” PLoS Pathog. 6: e1001193.

Hall, R. A. and N. A. R. Gow (2013). “Mannosylation in *Candida albicans:* role in cell wall function and immune recognition.” Molecular Microbiology.

Hao, R., M. Bondesson, A. V. Singh, A. Riu, C. W. McCollum, T. B. Knudsen, D. A. Gorelick and J.-Å. Gustafsson (2013). “Identification of estrogen target genes during zebrafish embryonic development through transcriptomic analysis.” PloS one 8: e79020–e79020.

Hopke, A., N. Nicke, E. E. Hidu, G. Degani, L. Popolo and R. T. Wheeler (2016). “Neutrophil Attack Triggers Extracellular Trap-Dependent *Candida* Cell Wall Remodeling and Altered Immune Recognition.” PLoS Pathogens 12: e1005644.

Inglis, D. O., M. B. Arnaud, J. Binkley, P. Shah, M. S. Skrzypek, F. Wymore, G. Binkley, S. R. Miyasato, M. Simison and G. Sherlock (2012). “The *Candida* genome database incorporates multiple *Candida* species: multispecies search and analysis tools with curated gene and protein information for *Candida albicans* and *Candida glabrata*.” Nucleic acids research 40: D667–D674.

Kenno, S., C. Speth, G. Rambach, U. Binder, S. Chatterjee, R. Caramalho, H. Haas, C. Lass Flörl, J. Shaughnessy and S. Ram (2018). “*Candida albicans* factor H binding molecule Hgt1p-a low glucose-induced transmembrane protein is trafficked to the cell wall and impairs phagocytosis and killing by human neutrophils.” Frontiers in microbiology 9: 3319.

Kinsman, O. S. and A. E. Collard (1986). “Hormonal factors in vaginal candidiasis in rats.” Infection and immunity 53: 498–504.

Klein, S. L. (2000). “The effects of hormones on sex differences in infection: from genes to behavior.” Neuroscience & Biobehavioral Reviews 24: 627–638.

Kozel, T. R., L. C. Weinhold and D. M. Lupan (1996). “Distinct characteristics of initiation of the classical and alternative complement pathways by *Candida albicans*.” Infect Immun 64: 3360–3368.

Lambris, J. D., D. Ricklin and B. V. Geisbrecht (2008). “Complement evasion by human pathogens.” Nature Reviews Microbiology 6: 132–142.

Lee, Y. J., G. R. Jeschke, F. M. Roelants, J. Thorner and B. E. Turk (2012). “Reciprocal phosphorylation of yeast glycerol-3-phosphate dehydrogenases in adaptation to distinct types of stress.” Molecular and cellular biology 32: 4705–4717.

Lopes, J. P., M. Stylianou, E. Backman, S. Holmberg, J. Jass, R. Claesson and C. F. Urban (2018). “Evasion of Immune Surveillance in Low Oxygen Environments Enhances *Candida albicans* Virulence.” mBio 9.

Luo, S., A. Hartmann, H.-M. Dahse, C. Skerka and P. F. Zipfel (2010). “Secreted pH-regulated antigen 1 of *Candida albicans* blocks activation and conversion of complement C3.” The Journal of Immunology 185: 2164–2173.

Luo, S., R. Hoffmann, C. Skerka and P. F. Zipfel (2012). “Glycerol-3-Phosphate Dehydrogenase 2 Is a Novel Factor H–, Factor H–like Protein 1–, and Plasminogen-Binding Surface Protein of *Candida albicans*.” The Journal of Infectious Diseases 207: 594–603.

Madani, N. D., P. J. Malloy, P. Rodriguez-Pombo, A. V. Krishnan and D. Feldman (1994). “*Candida albicans* estrogen-binding protein gene encodes an oxidoreductase that is inhibited by estradiol.” PNAS 91: 922–926.

Mallick, E. M., A. C. Bergeron, S. K. Jones, Z. R. Newman, K. M. Brothers, R. Creton, R. T. Wheeler and R. J. Bennett (2016). “Phenotypic Plasticity Regulates *Candida albicans* Interactions and Virulence in the Vertebrate Host.” Frontiers in Microbiology 7.

Marín, E., C. M. Parra-Giraldo, C. Hernández-Haro, M. L. Hernáez, C. Nombela, L. Monteoliva and C. Gil (2015). “*Candida albicans* Shaving to Profile Human Serum Proteins on Hyphal Surface.” Frontiers in microbiology 6: 1343–1343.

McClelland, E. E. and J. M. Smith (2011). “Gender specific differences in the immune response to infection.” Arch Immunol Ther Exp (Warsz) 59: 203–213.

Meri, T., H. Amdahl, M. J. Lehtinen, S. Hyvärinen, J. V. McDowell, A. Bhattacharjee, S. Meri, R. Marconi, A. Goldman and T. S. Jokiranta (2013). “Microbes Bind Complement Inhibitor Factor H via a Common Site.” PLOS Pathogens 9: e1003308.

Meri, T., A. Hartmann, D. Lenk, R. Eck, R. Würzner, J. Hellwage, S. Meri and P. F. Zipfel (2002). “The yeast *Candida albicans* binds complement regulators factor H and FHL-1.” Infection and immunity 70: 5185–5192.

Merle, N. S., R. Noe, L. Halbwachs-Mecarelli, V. Fremeaux-Bacchi and L. T. Roumenina (2015). “Complement System Part II: Role in Immunity.” Frontiers in immunology 6: 257–257.

Monteiro, P. T., J. Oliveira, P. Pais, M. Antunes, M. Palma, M. Cavalheiro, M. Galocha, C. P. Godinho, L. C. Martins, N. Bourbon, et al. (2019). “YEASTRACT+: a portal for cross-species comparative genomics of transcription regulation in yeasts.” Nucleic Acids Research 48: D642–D649.

Netea, M. G., G. D. Brown, B. J. Kullberg and N. A. R. Gow (2008). “An integrated model of the recognition of *Candida albicans* by the innate immune system.” Nature Reviews: Microbiology 6: 67–78.

Ngampasutadol, J., S. Ram, S. Gulati, S. Agarwal, C. Li, A. Visintin, B. Monks, G. Madico and P. A. Rice (2008). “Human Factor H Interacts Selectively with *Neisseria gonorrhoeae* and Results in Species-Specific Complement Evasion.” The Journal of Immunology 180: 3426.

Nobile, C. J. and A. P. Mitchell (2005). “Regulation of Cell-Surface Genes and Biofilm Formation by the *C. albicans* Transcription Factor Bcr1p.” Current Biology 15: 1150–1155.

Noble, S. M., S. French, L. A. Kohn, V. Chen and A. D. Johnson (2010). “Systematic screens of a *Candida albicans* homozygous deletion library decouple morphogenetic switching and pathogenicity.” Nature genetics 42: 590–598.

Pericolini, E., S. Perito, A. Castagnoli, E. Gabrielli, A. Mencacci, E. Blasi, A. Vecchiarelli and R. T. Wheeler (2018). “Epitope unmasking in vulvovaginal candidiasis is associated with hyphal growth and neutrophilic infiltration.” PLoS One 13: e0201436.

Poltermann, S., A. Kunert, M. von der Heide, R. Eck, A. Hartmann and P. F. Zipfel (2007). “Gpm1p is a factor H-, FHL-1-, and plasminogen-binding surface protein of *Candida albicans*.” Journal of biological chemistry 282: 37537–37544.

Pradhan, A., G. M. Avelar, J. M. Bain, D. Childers, C. Pelletier, D. E. Larcombe, E. Shekhova, M. G. Netea, G. D. Brown, L. Erwig, et al. (2019). “Non-canonical signalling mediates changes in fungal cell wall PAMPs that drive immune evasion.” Nature communications 10: 5315–5315.

Pradhan, A., G. M. Avelar, J. M. Bain, D. S. Childers, D. E. Larcombe, M. G. Netea, E. Shekhova, C. A. Munro, G. D. Brown, L. P. Erwig, et al. (2018). “Hypoxia Promotes Immune Evasion by Triggering ß-Glucan Masking on the *Candida albicans* Cell Surface via Mitochondrial and cAMP-Protein Kinase A Signaling.” mBio 9: e01318–01318.

Salinas-Muñoz, L., R. Campos-Fernández, E. Mercader, I. Olivera-Valle, C. Fernández-Pacheco, L. Matilla, J. García-Bordas, J. C. Brazil, C. A. Parkos, F. Asensio, et al. (2018). “Estrogen Receptor-Alpha (ESR1) Governs the Lower Female Reproductive Tract Vulnerability to *Candida albicans*.” Frontiers in immunology 9: 1033–1033.

Sanglard, D., F. Ischer, M. Monod and J. Bille (1997). “Cloning of *Candida albicans* genes conferring resistance to azole antifungal agents: characterization of *CDR2*, a new multidrug ABC transporter gene.” Microbiology 143 (Pt 2): 405–416.

Sherrington, S. L., E. Sorsby, N. Mahtey, P. Kumwenda, M. D. Lenardon, I. Brown, E. R. Ballou, D. M. MacCallum and R. A. Hall (2017). “Adaptation of *Candida albicans* to environmental pH induces cell wall remodelling and enhances innate immune recognition.” PLOS Pathogens 13: e1006403.

Sobel, J. D. (1988). “Pathogenesis and Epidemiology of Vulvovaginal Candidiasis.” Annals of the New York Academy of Sciences 544: 547–557.

Sobel, J. D. (1997). “Vaginitis.” New England Journal Medicine 337: 1896–1903.

Souder, J. P. and D. A. Gorelick (2017). “Quantification of Estradiol Uptake in Zebrafish Embryos and Larvae.” Toxicological sciences : an official journal of the Society of Toxicology 158: 465–474.

Sun, G., H. Li, Y. Wang, B. Zhang and S. Zhang (2010). “Zebrafish complement factor H and its related genes: Identification, evolution, and expression.” Functional and Integrative Genomics 10: 577–587.

Taneja, V. (2018). “Sex Hormones Determine Immune Response.” Frontiers in Immunology 9.

Tripathi, A., E. Liverani, A. Tsygankov and S. Puri (2020). “Iron alters the cell wall composition and intracellular lactate to affect *Candida albicans* susceptibility to antifungals and host immune response.” J Biol Chem.

van der Maten, E., B. van den Broek, M. I. de Jonge, K. J. W. Rensen, M. J. Eleveld, A. L. Zomer, A. J. H. Cremers, G. Ferwerda, R. de Groot, J. D. Langereis, et al. (2018). “*Streptococcus pneumoniae* PspC Subgroup Prevalence in Invasive Disease and Differences in Contribution to Complement Evasion.” Infection and immunity 86: e00010–00018.

van Lunzen, J. and M. Altfeld (2014). “Sex differences in infectious diseases-common but neglected.” J Infect Dis 209 Suppl 3: S79–80.

Walther, A. and J. Wendland (2003). “An improved transformation protocol for the human fungal pathogen *Candida albicans*.” Curr. Genetics 42: 339–343.

Wheeler, R. T. and G. R. Fink (2006). “A Drug-Sensitive Genetic Network Masks Fungi from the Immune System.” PLoS Pathog 2: e35.

Wheeler, R. T., D. Kombe, S. D. Agarwala and G. R. Fink (2008). “Dynamic, Morphotype-Specific *Candida albicans* ß-Glucan Exposure during Infection and Drug Treatment.” PLoS Pathogens 4: e1000227.

White, S. and B. Larsen (1997). “*Candida albicans* morphogenesis is influenced by estrogen.” Cellular and Molecular Life Sciences CMLS 53: 744–749.

Williams, R. B. and M. C. Lorenz (2020). “Multiple Alternative Carbon Pathways Combine To Promote *Candida albicans* Stress Resistance, Immune Interactions, and Virulence.” mBio 11.

Wilson, R. B., D. Davis, B. M. Enloe and A. P. Mitchell (2000). “A recyclable *Candida albicans URA3* cassette for PCR product-directed gene disruptions.” Yeast 16: 65–70.

Zhang, S. and P. Cui (2014). “Complement system in zebrafish.” Developmental & Comparative Immunology 46: 3–10.

