## Supplemental figures and tables for "Oestrogen promotes innate immune evasion of *Candida albicans* through inactivation of the alternative complement system"

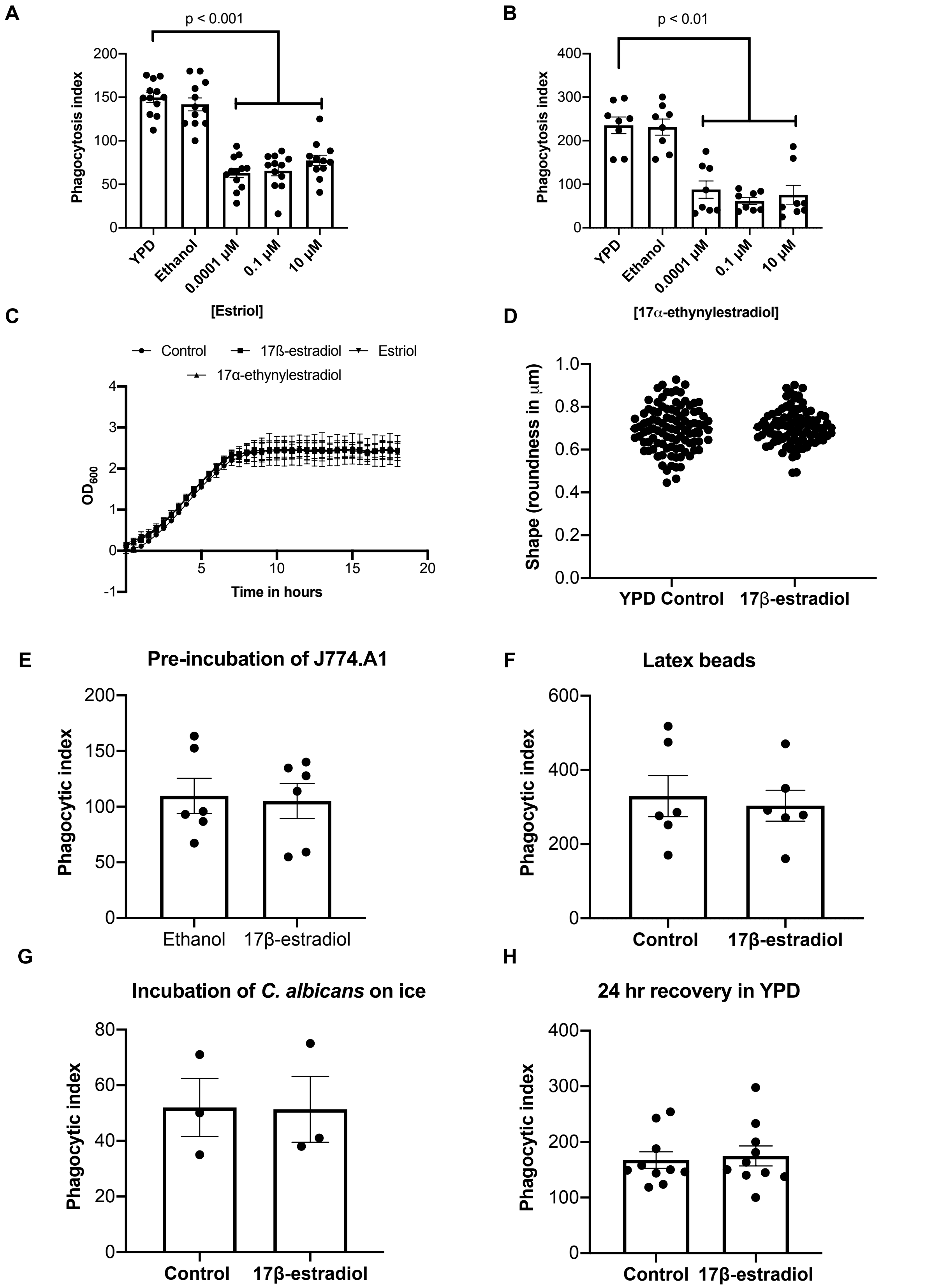

**Figure S1. Oestrogen inhibits *C. albicans* phagocytosis, but does not affect fungal growth or morphology. A)** J774A.1 phagocytosis rates of *C. albicans* (SC5314) grown in the presence of estriol. **B)** J774A.1 phagocytosis rates of *C. albicans* grown in the presence of 17α-ethynylestradiol. **C)** *C. albicans* cells were grown in 96-welled plate in YPD broth supplemented with 10 µM 17β-estradiol, 10 µM 17α-ethynylestradiol or 10 µM estriol. Optical densities (OD) of the cultures were recorded every 30 min for 18 h. **D)** *C. albicans* was grown in YPD broth supplemented with 17β-estradiol 10 µM for 4 h. Cells were washed with PBS, stained with concanavalin A, imaged by microscopy and analysed for shape. **E)** J774A.1 macrophages preincubated with 10 µM 17β-estradiol were infected with *C. albicans* cells grown in YPD and phagocytosis rates quantified. **F)** Latex beads were incubated at room temperature in PBS with or without 10 µM 17β-estradiol. Beads were washed and co-incubated with J774A.1 macrophages and phagocytosis rates quantified. **G)** *C. albicans* cells were incubated in PBS at 4°C for 4 h with or without 10 µM 17β-estradiol. Cells were washed and co-incubated with J774A.1 macrophages and phagocytosis rates quantified **H)** *C. albicans* cells previously grown in YPD with or without 10 µM 17β-estradiol were harvested, washed and re-incubated in fresh YPD for 24 h. Cells were co-incubated with J774A.1 macrophages and phagocytosis rates quantified. All data represent the mean ± SEM from at least three independent experiments.

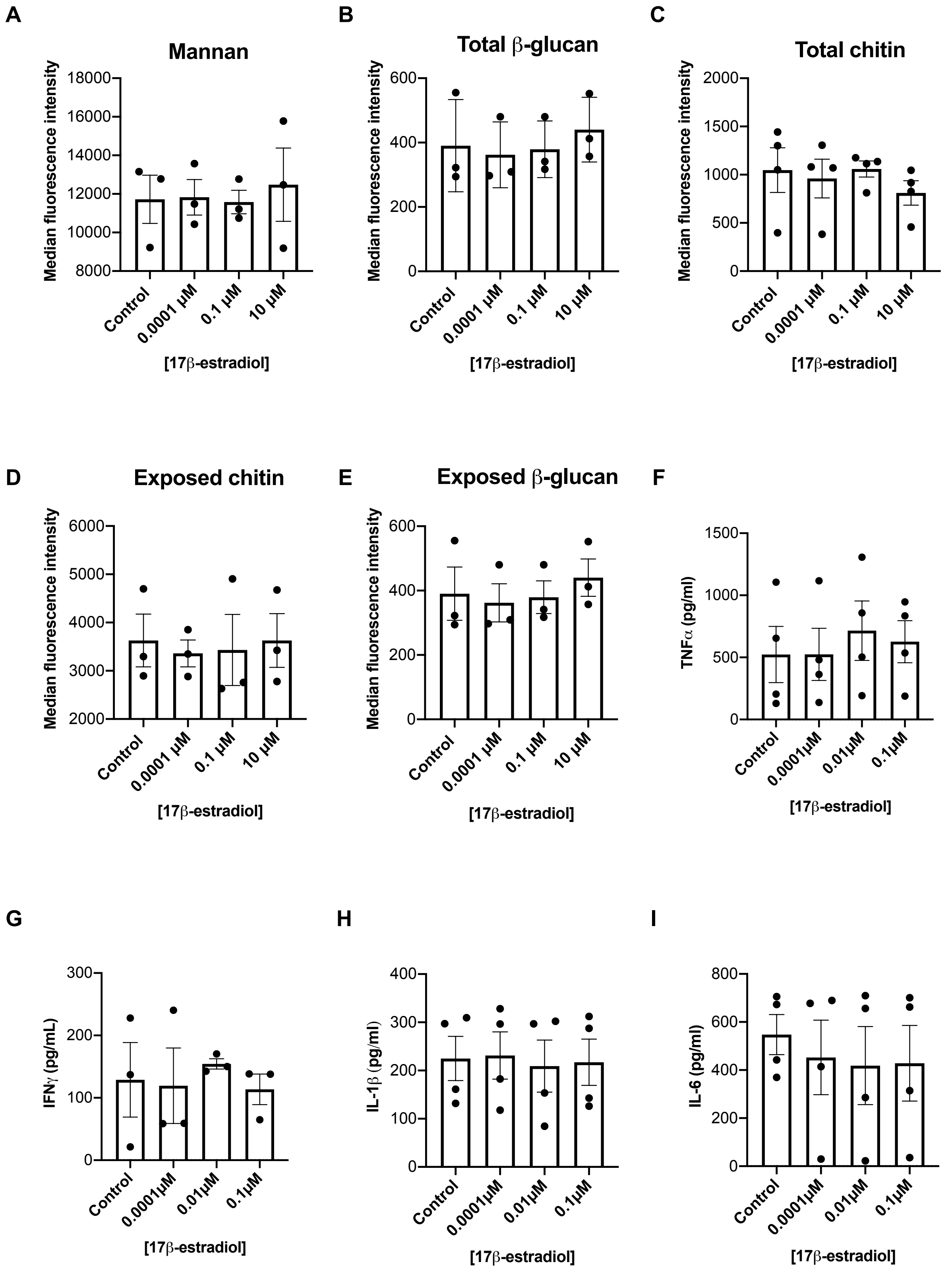

**Figure S2. Reduced phagocytosis rates are not correlated with altered cytokine secretion or gross changes in cell wall carbohydrates.** *C. albicans* cells were grown in YPD with or without 10 μM 17β-estradiol. Cells were harvested, washed in PBS, fixed with 4%

PFA and stained for **A**) total mannan **B**) total glucan **C**) total chitin **D)** exposed chitin **E)** exposed glucan. Staining was quantified by flow cytometry and median fluorescence intensities (MFI) determined. All data represent the mean ± SEM from at least three independent biological experiments. *C. albicans* cells were grown in YPD with or without 10 μM 17β-estradiol for 4 h. Cells were washed, fixed with 4% PFA and co-incubated with PBMCs for 24h. Secretion of **F**) TNFα, **G)** IFNγ, **H)** IL-1β and **I)** IL-6 by PBMCs was quantified by ELISA. Data represent the mean ± SEM from four independent experiments using different donors.

**Table S2. Gene Ontology analysis of differentially regulated genes**

| GO Term | Cluster | Adjusted P-value | Genes |
|---|---|---|---|
| **Up regulated genes** | | | |
| Hormone binding | 2 / 59 genes (3.4%) | 0.00473 | *CDR1, EBP1* |
| snoRNA binding | 4 / 59 genes (6.8%) | 0.00541 | *BMS1, NOP14, HCA4, NOP10* |
| catalytic activity, acting on a rRNA | 3 / 59 genes (5.1%) | 0.01756 | *DIM1, SPB1, CR_04170W_A* |
| rRNA methyltransferase activity | 3 / 59 genes (5.1%) | 0.01756 | *DIM1, SPB1, CR_04170W_A* |
| U3 snoRNA binding | 3 / 59 genes (5.1%) | 0.02147 | *BMS1, NOP14, HCA4* |
| FMN binding | 3 / 59 genes (5.1%) | 0.0737 | *OYE32, EBP1, OYE23* |
| **Down regulated genes** | | | |
| Oxidoreductase activity | 19 / 76 genes (25.0%) | 2.07E-05 | *HMX1, CSH1, IFD6, C1_04460C_A, CAT1, GCV2, C2_00180C_A, ALD5, C2_04480W_A, PST1, PST2, FOX2, POX1-3, MET13, SOU1, GRP2, HPD1, FDH1, FRE7* |
| Oxidoreductase activity, acting on the CH-OH group of donors, NAD or NADP as acceptor | 7 / 76 genes (9.2%) | 0.00078 | *CSH1, IFD6, FOX2, SOU1, GRP2, HPD1, FDH1* |
| Oxidoreductase activity, acting on CH-OH group of donors | 7 / 76 genes (9.2%) | 0.00115 | *CSH1, IFD6, FOX2, SOU1, GRP2, HPD1, FDH1* |

| | | | |
|---|---|---|---|
| Aryl-alcohol dehydrogenase (NAD+) activity | 2 / 76 genes (2.6%) | 0.02754 | *CSH1, IFD6* |
| NAD(P)H dehydrogenase (quinone) activity | 2 / 76 genes (2.6%) | 0.05466 | *PST1, PST2* |
| Glutaminase activity | 2 / 76 genes (2.6%) | 0.05466 | *SNZ1, SNO1* |
| Glucose transmembrane transporter activity | 3 / 76 genes (3.9%) | 0.08953 | *HGT19, HGT5, HGT10* |

**Table S3. Motif letter-probability matrix**

| *Motif* | *A* | *C* | *G* | *T* |
|---|---|---|---|---|
| ***Motif 1*** ***E-value = 1.2e-159*** | | | | |
| | 0.000000 | 0.037037 | 0.175926 | 0.787037 |
| | 0.009259 | 0.046296 | 0.129630 | 0.814815 |
| | 0.000000 | 0.379630 | 0.000000 | 0.620370 |
| | 0.101852 | 0.074074 | 0.064815 | 0.759259 |
| | 0.000000 | 0.120370 | 0.055556 | 0.824074 |
| | 0.018519 | 0.259259 | 0.009259 | 0.712963 |
| | 0.000000 | 0.259259 | 0.000000 | 0.740741 |
| | 0.055556 | 0.129630 | 0.009259 | 0.805556 |
| | 0.000000 | 0.240741 | 0.009259 | 0.750000 |
| | 0.000000 | 0.157407 | 0.129630 | 0.712963 |
| | 0.083333 | 0.416667 | 0.027778 | 0.472222 |
| | 0.151389 | 0.232870 | 0.001389 | 0.614352 |
| | 0.000000 | 0.185185 | 0.000000 | 0.814815 |
| | 0.055556 | 0.129630 | 0.046296 | 0.768519 |
| | 0.018519 | 0.546296 | 0.120370 | 0.314815 |
| | 0.083333 | 0.037037 | 0.018519 | 0.861111 |
| | 0.037037 | 0.157407 | 0.092593 | 0.712963 |
| | 0.129630 | 0.277778 | 0.046296 | 0.546296 |
| | 0.166667 | 0.314815 | 0.055556 | 0.462963 |
| | 0.129630 | 0.092593 | 0.037037 | 0.740741 |
| | 0.138889 | 0.222222 | 0.083333 | 0.555556 |
| | 0.129630 | 0.157407 | 0.055556 | 0.657407 |
| | 0.037037 | 0.194444 | 0.037037 | 0.731481 |
| | 0.194444 | 0.000000 | 0.111111 | 0.694444 |
| | 0.083333 | 0.333333 | 0.120370 | 0.462963 |
| | 0.203704 | 0.268519 | 0.027778 | 0.500000 |
| | 0.185185 | 0.342593 | 0.046296 | 0.425926 |
| | 0.175926 | 0.194444 | 0.037037 | 0.592593 |
| | 0.111111 | 0.287037 | 0.101852 | 0.500000 |
| ***Motif 2*** ***E-value = 4.9e-04*** | | | | |
| | 0.321429 | 0.392857 | 0.071429 | 0.214286 |
| | 0.232143 | 0.214286 | 0.285714 | 0.267857 |
| | 0.357143 | 0.375000 | 0.000000 | 0.267857 |

| | | | | |
|---|---|---|---|---|
| | 0.071429 | 0.464286 | 0.000000 | 0.464286 |
| | 0.125000 | 0.428571 | 0.107143 | 0.339286 |
| | 0.107143 | 0.178571 | 0.000000 | 0.714286 |
| | 0.232143 | 0.464286 | 0.017857 | 0.285714 |
| | 0.089286 | 0.392857 | 0.053571 | 0.464286 |
| | 0.178571 | 0.678571 | 0.125000 | 0.017857 |
| | 0.166964 | 0.466964 | 0.002679 | 0.363393 |
| | 0.000000 | 0.821429 | 0.053571 | 0.125000 |
| | 0.232143 | 0.375000 | 0.017857 | 0.375000 |
| | 0.000000 | 0.660714 | 0.000000 | 0.339286 |
| | 0.000000 | 0.732143 | 0.000000 | 0.267857 |
| | 0.196429 | 0.482143 | 0.000000 | 0.321429 |
| | 0.017857 | 0.660714 | 0.089286 | 0.232143 |
| | 0.107143 | 0.589286 | 0.035714 | 0.267857 |
| | 0.000000 | 0.589286 | 0.000000 | 0.410714 |
| | 0.000000 | 0.821429 | 0.000000 | 0.178571 |
| | 0.375000 | 0.089286 | 0.053571 | 0.482143 |
| | 0.392857 | 0.482143 | 0.125000 | 0.000000 |
| ***Motif 3*** ***E-value = 2.5e-009*** | | | | |
| | 0.805556 | 0.055556 | 0.138889 | 0.000000 |
| | 0.763889 | 0.097222 | 0.000000 | 0.138889 |
| | 0.500000 | 0.430556 | 0.069444 | 0.000000 |
| | 0.722222 | 0.069444 | 0.111111 | 0.097222 |
| | 0.902778 | 0.013889 | 0.083333 | 0.000000 |
| | 0.652778 | 0.222222 | 0.000000 | 0.125000 |
| | 0.847222 | 0.000000 | 0.152778 | 0.000000 |
| | 0.972222 | 0.027778 | 0.000000 | 0.000000 |
| | 0.847222 | 0.083333 | 0.069444 | 0.000000 |
| | 0.902778 | 0.027778 | 0.000000 | 0.069444 |
| | 1.000000 | 0.000000 | 0.000000 | 0.000000 |
| | 0.791667 | 0.208333 | 0.000000 | 0.000000 |
| | 1.000000 | 0.000000 | 0.000000 | 0.000000 |
| | 1.000000 | 0.000000 | 0.000000 | 0.000000 |

**Table S4. Transcription factors that bind identified DNA binding motifs**

| Motif | *S. cerevisiae* | *C. albicans* | Description on CGD |
|---|---|---|---|
| **Motif 1** | Azf1 | CR_02510W | Induced by Mnl1 under weak acid stress |
| | Fkh1 | Fkh2 | Morphogenesis regulator |
| | Hsf1 | Cta8 | Mediates heat shock transcriptional induction |
| | Sfl1 | Sfl1 | Negative regulation of morphogenesis, flocculation and virulence |

| | | | |
|---|---|---|---|
| | Ste12 | Cph1 | Mating, and filamentation on solid media repressed |
| | Cup2 | NA | |
| | Fkh2 | NA | |
| **Motif 2** | Haa1 | Cup2 | Required for normal resistance to copper |
| | Msn2 | Msn4 | Similar to *S. cerevisiae* Msn4 |
| | Msn4 | Msn4 | Similar to *S. cerevisiae* Msn4 |
| | Ygr067c | Try5 | Regulator of yeast form adherence |
| | Rap1 | Rap1 | Binds telomeres and regulatory sequences in DNA |
| | Yml081w | Zms1 | Spider biofilm induced |
| | Cha4 | Tea1 | Putative transcription factor with zinc cluster DNA-binding motif |
| | Nrg1 | Nrg1 | Regulates chlamydospore formation, hyphal gene induction, virulence |
| | Rgm1 | NA | |
| | Rph1 | Rph1 | Ortholog(s) have DNA-binding transcription repressor activity |
| | Gis1 | NA | |
| | Usv1 | Bcr1 | Regulates a/alpha biofilm formation, matrix, cell-surface-associated genes |
| | Yer130c | Mnl1 | induces transcripts of stress response genes via SLE (STRE-like) elements |
| | Ypr022c | C3_06150W | Ortholog(s) have role in negative regulation of transcription by RNA polymerase II |
| **Motif 3** | Azf1 | CR_02510W | Induced by Mnl1 under weak acid stress |
| | Cup2 | NA | |
| | Hcm1 | Hcm1 | Similar to *S. cerevisiae* Hcm1 |
| | Fkh2 | NA | |

**Table S5. GO Ter analysis of *BCR1* regulated gene**

| **GO ID** | **GO Term** | **% of input genes** | **p-value** | **Genes** |
|---|---|---|---|---|
| GO:0009986 | cell surface | 15.56% | 4.91E-07 | *ALS4, ECM331, GPD2, PGA13, RBT5, SAP10, SIM1* |
| GO:0005886 | plasma membrane | 15.56% | 4.61E-05 | *AQY1, CFL5, DIP5, ECM331, GPR1, JEN1, RBT5* |
| GO:0009277 | fungal-type cell wall | 13.33% | 6.29E-08 | *ALS4, ECM331, PGA13, RBT5, SAP10, SIM1* |

| | | | | |
|---|---|---|---|---|
| GO:0016021 | integral component of membrane | 13.33% | 9.06E-05 | C1_00660C_A, *GPR1, JEN1, MAL31, RBD1, UFE1* |
| GO:0005634 | nucleus | 13.33% | 1.81E-03 | CR_03700C_A, *GRF10, SEF2, SFL2, WOR2, WOR3* |

**Table S6. Strains used in this study**

| Strain | Genotype | Source/reference |
|---|---|---|
| SC5314 | Wild type | (Gillum *et al.* 1984) |
| SVS006B | Clinical isolate | Prof Ramage, Glasgow University |
| SVS062A | Clinical isolate | Prof Ramage, Glasgow University |
| SN152 | *arg4Δ/arg4Δ leu2Δ/leu2Δ his1Δ/his1Δ URA3/ura3Δ:: λimm434 IRO1/iro1Δ:: λimm434* | (Noble *et al.* 2010) |
| SN250 | *his1Δ/his1Δ, leu2Δ::C. dubliniensis HIS1 /leu2Δ::C. maltosa LEU2-arg4Δ/arg4Δ, URA3/ura3Δ::imm434-IRO1/iro1Δ::imm434* | (Noble *et al.* 2010) |
| SN250-CIP30 | As SN250 but *RPS1/rps1::*CIP30 | This study |
| *rob1Δ* | As SN152 but *rob1Δ::C. dubliniensisHIS1/rob1Δ::C. maltose LEU2* | (Noble *et al.* 2010) |
| *bcr1Δ* | *arg4Δ/arg4Δ leu2Δ/leu2Δ his1Δ/his1Δ URA3/ura3Δ::λimm434 IRO1/iro1Δ:: λimm434 bcr1::LEU/bcr1::HIS1* | (Noble *et al.* 2010) |
| *gpd2Δ* | As SN152 but *gpd2Δ::C. dubliniensisHIS1/gpd2Δ::C. maltose LEU2* | (Noble *et al.* 2010) |

| | | |
|---|---|---|
| *gpd2Δ*–CIP30 | As SN152 but *gpd2Δ::C. dubliniensisHIS1/gpd2Δ::C. maltose LEU2, RPS1/rps1::CIP30* | This study |
| *gpd2Δ*-CIP30-*GPD2* | As SN152 but *gpd2Δ::C. dubliniensisHIS1/gpd2Δ::C. maltose LEU2RPS1/rps1::CIP30-GPD2* | This study |
| CAI4 | *ura3::imm434/ura3::imm434 iro1/iro1::imm434* | (Fonzi and Irwin 1993) |
| CAI4-pSM2 | *ura3::imm434/ura3::imm434 iro1/iro1::imm434::*pSM2 | (Hall *et al.* 2010) |
| CAI4-pSM2-*GPD2* | *ura3::imm434/ura3::imm434 iro1/iro1::imm434::*pSM2-pTef2-*GPD2* | This study |
| *ebp1*Δ | As CAI4 but *ebp1::*dlp200/*ebp1::*dlp200 | This study |
| *ebp1*Δ-pSM2 | As CAI4 but *ebp1::*dlp200/*ebp1*::dlp200-pSM2 | This study |
| CAF2-1 | *URA3/ura3:: λimm434* | (Fonzi and Irwin 1993) |
| *cdr1*Δ | As CAF2-1, but *crd1::hisG/cdr1:hisG-URA3-HisG* | (Sanglard *et al.* 1997) |
| *cdr1/2*Δ | As CAF2-1, but *crd1::hisg/cdr1:hisG, cdr2::hisG-URA3-hisG/cdr2::hisG* | (Sanglard *et al.* 1997) |

**Table S7. Primers used in this study**

| **Primer** | **Sequence** |
|---|---|
| GPD2-SacI-F | CTCCGAGCTCGGTGATGGTGATGGTGATGG |
| GPD2-NotI-R | GGAGAGCGGCCGCTGGTAAATTGGACAACGAGTGG |
| EBP1-5F: | CTCCGATATCATCGCATGAG |
| EBP1-5R: | GGAGAGGAGCTCctgatgatatgataatttgc |
| EBP1-3F: | GGAGAGAAGCTTGGGAATGAAGTTCTATTAGC |

| | |
|---|---|
| EBP1-3R: | GGAGAGGGTACCgttacatctactactacagg |
| RT-ACT1-F | CCTACGTGTACTTGTGCAAGGCAA |
| RT-ACT1-R | TAGTTGTGTGCACTGAGCGTCGAA |
| RT-GDP2-F | GCCAACGAAGTTGCCAAAGGT |
| RT-GPD2-R | AGGCACCAGCAATAGAGGCA |
